# A Specific Neuroligin3-αNeurexin1 Code Regulates GABAergic Synaptic Function in Mouse Hippocampus

**DOI:** 10.1101/2020.05.27.119024

**Authors:** Motokazu Uchigashima, Kohtarou Konno, Emily Demchak, Amy Cheung, Takuya Watanabe, David Keener, Manabu Abe, Timmy Le, Kenji Sakimura, Toshikuni Sasaoka, Takeshi Uemura, Yuka Imamura Kawasawa, Masahiko Watanabe, Kensuke Futai

**Affiliations:** Brudnick Neuropsychiatric Research Institute, Department of Neurobiology, University of Massachusetts Medical School, 364 Plantation Street, LRB-706 Worcester, MA 01605-2324, USA; Department of Cellular Neuropathology, Brain Research Institute, Niigata University, Niigata 951-8585, Japan; Department of Anatomy, Faculty of Medicine, Hokkaido University, Sapporo, Hokkaido 060-8638, Japan; Department of Biochemistry and Molecular Biology and Institute for Personalized Medicine, Pennsylvania State University College of Medicine, 500 University Drive, Hershey, Pennsylvania 17033, USA; Department of Animal Model Development, Brain Research Institute, Niigata University, Niigata 951-8585, Japan; Department of Comparative and Experimental Medicine, Brain Research Institute, Niigata University, Niigata 951-8585, Japan; Division of Gene Research, Research Center for Supports to Advanced Science, Shinshu University, Nagano 390-8621, Japan; Institute for Biomedical Sciences, Interdisciplinary Cluster for Cutting Edge Research, Shinshu University, Nagano 390-8621, Japan; Department of Pharmacology Pennsylvania State University College of Medicine, 500 University Drive, Hershey, Pennsylvania 17033, USA

**Keywords:** neuroligin, neurexin, cholecystokinin, vesicular glutamate transporter 3, trans-synaptic interaction, GABAergic synapses, hippocampus, mouse

## Abstract

Synapse formation and regulation require interactions between pre- and postsynaptic proteins, notably cell adhesion molecules (CAMs). It has been proposed that the functions of neuroligins (Nlgns), postsynaptic CAMs, rely on the formation of trans-synaptic complexes with neurexins (Nrxns), presynaptic CAMs. Nlgn3 is a unique Nlgn isoform that localizes at both excitatory and inhibitory synapses. However, Nlgn3 function mediated via Nrxn interactions is unknown. Here, we demonstrate that Nlgn3 localizes at postsynaptic sites apposing vesicular glutamate transporter 3-expressing (VGT3+) inhibitory terminals and regulates VGT3+ inhibitory interneuron-mediated synaptic transmission in mouse organotypic slice cultures. Gene expression analysis of interneurons revealed that the αNrxn1+AS4 splice isoform is highly expressed in VGT3+ interneurons as compared with other interneurons. Most importantly, postsynaptic Nlgn3 requires presynaptic αNrxn1+AS4 expressed in VGT3+ interneurons to regulate inhibitory synaptic transmission. Our results indicate that specific Nlgn-Nrxn interactions generate distinct functional properties at synapses.

## Introduction

In central synapses, cell adhesion molecules (CAMs) are major players in trans-synaptic interactions (de Wit & Ghosh, 2016) which serve a primary role in initiating synapse formation by directing contact between axonal and dendritic membranes. Emerging evidence suggests that trans-synaptic interactions are also important for synapse identity, function, plasticity and maintenance (Biederer, Kaeser, & Blanpied, 2017; Campbell & Tyagarajan, 2019; Sudhof, 2017). Numerous CAM variants exist due to large gene families and alternative splicing, generating a vast array of possible combinations of pre- and postsynaptic CAMs. Although some specific trans-synaptic interactions of CAMs have been reported to underlie distinct synaptic properties (Chih, Gollan, & Scheiffele, 2006; Fossati et al., 2019; Futai, Doty, Baek, Ryu, & Sheng, 2013), elucidating synaptic CAM complexes that dictate synapse identity and function remains a major challenge.

Four neuroligin (Nlgn) genes (*Nlgn1, Nlgn2, Nlgn3* and *Nlgn4*) encode postsynaptic CAMs (Nlgn1, Nlgn2, Nlgn3 and Nlgn4) that contain extracellular cholinesterase-like domains and transmembrane and PDZ-binding motif-containing intracellular domains. Each Nlgn protein has a distinct pattern of subcellular localization at excitatory, inhibitory, dopaminergic and cholinergic synapses (Song, Ichtchenko, Sudhof, & Brose, 1999; Takacs, Freund, & Nyiri, 2013; Uchigashima, Ohtsuka, Kobayashi, & Watanabe, 2016; Varoqueaux, Jamain, & Brose, 2004). Interestingly, Nlgn3 is the only Nlgn isoform localized at both excitatory and inhibitory synapses (Baudouin et al., 2012; Budreck & Scheiffele, 2007; Uchigashima et al., 2020), regulating their synaptic functions (Etherton et al., 2011; Foldy, Malenka, & Sudhof, 2013; Horn & Nicoll, 2018; Shipman et al., 2011; Tabuchi et al., 2007). However, the trans-synaptic framework that dictates Nlgn3 function is poorly understood.

Neurexins (Nrxns) are presynaptic CAMs produced from three genes (*Nrxn1, Nrxn2, Nrxn3*) that are transcribed from different promoters as longer alpha (*αNrxn1* - *3*), shorter beta (*βNrxn1* - *3*) and *Nrxn1*-specific gamma (*γNrxn1*) isoforms (Sterky et al., 2017; Tabuchi & Sudhof, 2002), and serve as the sole presynaptic binding partners for Nlgns. Each *Nrxn* gene has six alternative splicing sites, named AS1 - AS6, resulting in thousands of potential *Nrxn* splice isoforms (Gorecki, Szklarczyk, Lukasiuk, Kaczmarek, & Simons, 1999; Missler, Fernandez-Chacon, & Sudhof, 1998; Puschel & Betz, 1995; Schreiner et al., 2014; Treutlein, Gokce, Quake, & Sudhof, 2014; Ullrich, Ushkaryov, & Sudhof, 1995). Unique transcription patterns of *Nrxns* have been observed in hippocampal interneurons, suggesting that Nrxn proteins may determine the properties of GABAergic synapses in an input cell-dependent manner (Fuccillo et al., 2015).

Nrxn-Nlgn interactions depend on Nrxn protein length (long form [α] vs short form [β]), splice insertions at AS4 of Nrxns and splice insertions of Nlgns. For example, Nlgn1 splice variants that have splice insertions at site B have higher binding affinities for βNrxn1-AS4 (βNrxn1 lacking alternative splice insertion at AS4) than for βNrxn1+AS4 (containing an alternative splice insertion at AS4) (Boucard, Chubykin, Comoletti, Taylor, & Sudhof, 2005; Koehnke et al., 2010; Reissner, Klose, Fairless, & Missler, 2008). However, it is largely unknown which Nrxn-Nlgn combination defines specific synapse functionality. We recently found that Nlgn3Δ, which lacks both of the A1 and A2 alternative splice insertions, is the major Nlgn3 splice isoform expressed in hippocampal CA1 pyramidal neurons and regulates both excitatory and inhibitory synaptic transmission (Uchigashima et al., 2020). However, to the best of our knowledge, the synapses at which Nlgn3Δ interacts with presynaptic Nrxn isoform(s) have not been identified.

Interneurons exhibit extraordinary morphological, physiological and molecular diversity in the cortex and hippocampus (Klausberger & Somogyi, 2008; Markram et al., 2004; Pelkey et al., 2017; P. Somogyi & Klausberger, 2005). Indeed, there are over 20 classes of inhibitory interneurons in the CA1 area based on molecular markers, action potential firing patterns and morphology (Klausberger & Somogyi, 2008; Pelkey et al., 2017). Among them, interneurons expressing parvalbumin (Pv+), somatostatin (Sst+) and cholecystokinin (Cck+) display different morphologies, excitability and synaptic functions. However, the molecular mechanisms underlying their diversity are unknown.

In the present study, we show that Nlgn3 is selectively enriched at vesicular glutamate transporter 3-expressing (VGT3+) Cck+ inhibitory terminals in the hippocampal CA1 area. Gain-of-function and loss-of-function studies revealed that Nlgn3 regulates VGT3+ interneuron-mediated inhibitory synaptic transmission. Importantly, the effect of Nlgn3 on VGT3+ synapses was hampered by the deletion of all *Nrxn* genes in VGT3+ interneurons and rescued by the selective expression of αNrxn1+AS4 in VGT3+ interneurons. These results suggest that the trans-synaptic interaction between αNrxn1+AS4 and Nlgn3 underlies the input cell-dependent control of VGT3+ GABAergic synapses in the hippocampus.

## Results

### Nlgn3 is enriched at VGT3+ GABAergic synapses in the hippocampal CA1 region

In a recent study, we demonstrated that Nlgn3 localizes at and regulates both inhibitory and excitatory synapses in the hippocampal CA1 area (Uchigashima et al., 2020). However, the distribution of Nlgn3 at different types of inhibitory synapses has not yet been addressed. Therefore, we first examined which GABAergic inhibitory synapses express Nlgn3 in the CA1 area by immunohistochemistry. Cck+, Pv+ and Sst+ interneurons are the primary inhibitory neurons in the hippocampus. Moreover, the cell bodies and dendritic shafts of CA1 pyramidal cells are targeted by Cck+ and Pv+, and Sst+ interneurons, respectively (Pelkey et al., 2017). Our Nlgn3 antibody with specific immunoreactivity was validated in Nlgn3 KO brain (**Figure 1A**) and displayed a typical membrane protein distribution pattern in the hippocampus, as we recently reported (**Figure 1B, C**) (Uchigashima et al., 2020). Inhibitory synapses expressing Nlgn3 were identified by co-localization of Nlgn3 signals with vesicular inhibitory amino acid transporter (VIAAT) signals. Four different inhibitory axons/terminals were visualized by anti-VGT3 and -CB1 (markers for Cck+ interneurons) (Fruh et al., 2016), -Pv and -Sst antibodies. We found that punctate signals for Nlgn3 were frequently associated with GABAergic terminals co-labeled with VGT3 or CB1 (**Figure 1D-G and L**) compared with Pv (**Figure 1H, I**, and **L**) or Sst (**Figure 1J, K and L**) in the CA1 pyramidal cell layer. These data strongly suggest that Nlgn3 is preferentially recruited to Cck+ GABAergic synapses, but not to Pv+ or Sst+ inhibitory synapses.

**Figure 1.**
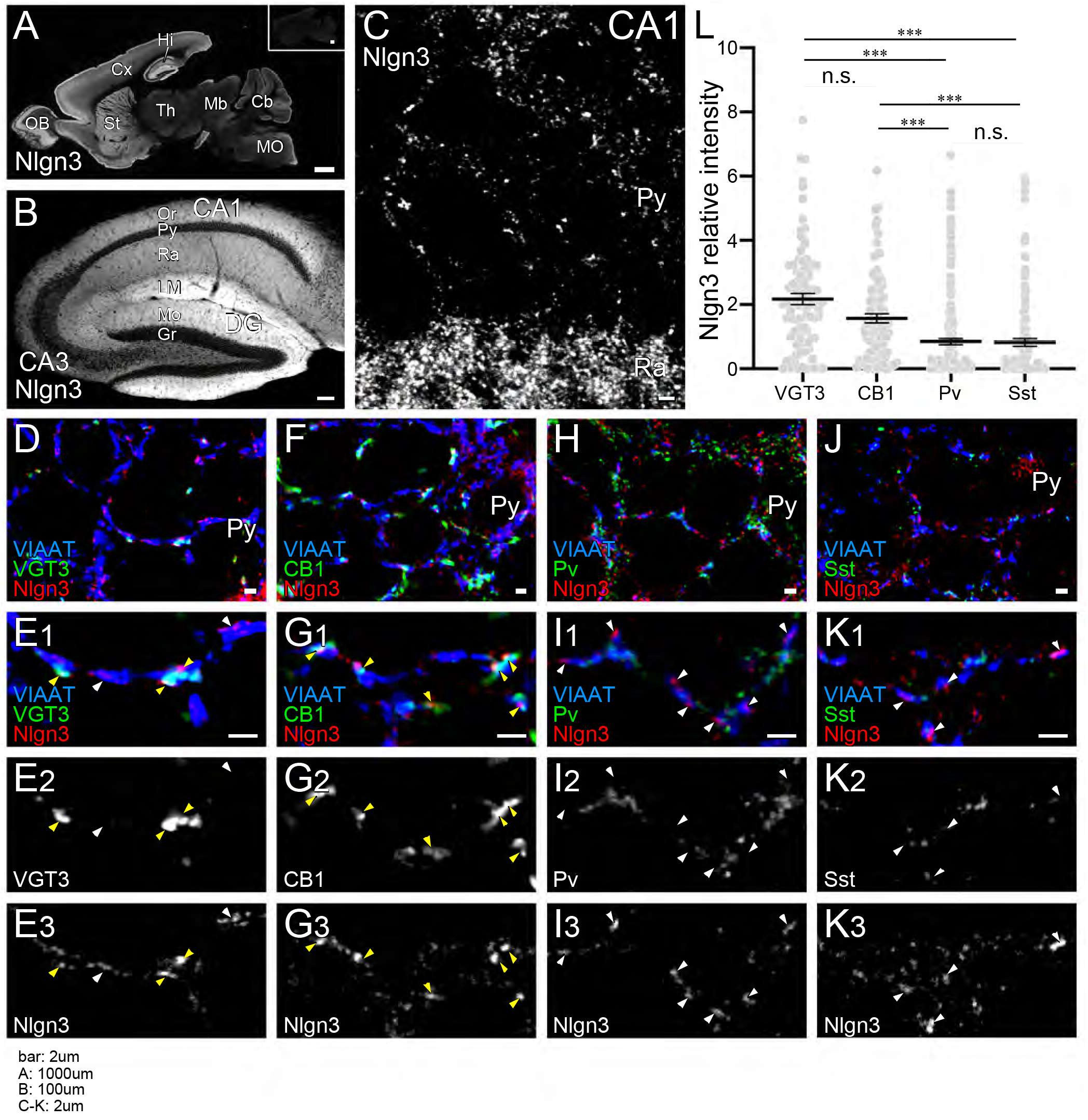
Nlgn3 is predominantly expressed in VGT3+ and CB1+ inhibitory synapses. **(A)** Immunofluorescence for Nlgn3 shows wide but distinct regional distribution in sagittal sections of P21 mouse brain. Higher levels were detected in the cortex (**A**, Cx), striatum (**A**, St), olfactory bulb (**A**, OB) and hippocampal formation (**A, B**, Hi). This staining was not detected in Nlgn3 KO mice (**A** inset). In the hippocampus, intense labeling was found in the st. lacunosum-moleculare (LM) of Ammon’s horn and in the molecular layer of dentate gyrus **(B)**. At high magnifications, Nlgn3 formed basket-like clustering around the somata of pyramidal cells in the CA1 region. **(D-K)** Triple immunofluorescence for Nlgn3, VIAAT and inhibitory terminal markers: VGT3 **(D, E)**, CB1 **(F, G)**, Pv **(H, I)** and Sst **(J, K)**. **(L)** Summary of the relative intensity of immunofluorescent signals for Nlgn3 at different inhibitory synapses. ***p < 0.001; n.s. not significant; Mann–Whitney U test. Bars on each column represent mean ± SEM. CA1–3, CA1–3 regions of the Ammon’s horn; Cb, cerebellum; Cx, cerebral cortex; DG, dentate gyrus; Gr, granule cell layer; Hi, hippocampus; LM, st. lacunosum-moleculare; Mb, midbrain; Mo, molecular layer; MO, medulla oblongata; OB, olfactory bulb; Or, st. oriens; Py, pyramidal cell layer; Ra, st. radiatum; St, striatum; Th, thalamus; Scale bars, 1000 μm **(A)**, 100 μm **(B)**, 2 μm **(C-K)**.

### Nlgn3 regulates inhibitory synaptic transmission at VGT3+ GABAergic synapses

To determine whether Nlgn3 has specific roles at VGT3+ inhibitory synapses, we assessed the effect of overexpressing Nlgn3Δ, the major Nlgn3 splice isoform expressed in CA1 pyramidal neurons (Uchigashima et al., 2020), on input-specific inhibitory transmission. To distinguish a subset of GABAergic synapses and evoke cell-specific synaptic transmission, we generated three cell type-specific fluorescent lines by crossing VGT3^cre^, Sst^cre^ or Pv^cre^ with a TdTomato (RFP) reporter line, producing respectively, VGT3/RFP, Sst/RFP and Pv/RFP mouse lines. TdTomato-expressing cells in each of the three fluorescent mouse lines were distributed in the CA1 in a layer-dependent manner (**Figure S1**). We evaluated the effect of Nlgn3Δ overexpression (OE) on unitary inhibitory postsynaptic currents (uIPSCs) by triple whole-cell recordings using organotypic slice cultures from each mouse line. Two to three days after transfection of *Nlgn3Δ* or EGFP control by biolistic gene gun, current- and voltage-clamp recordings were conducted from a presynaptic RFP+ interneuron and postsynaptic EGFP or EGFP/*Nlgn3Δ*-positive and -negative postsynaptic pyramidal neurons, respectively (**Figure 2A**). RFP interneurons expressing VGT3 in the pyramidal cell layer and stratum (st.) radiatum, Pv in the st. pyramidale and Sst in the st. oriens, were chosen (**Figure S1**). Unitary inhibitory postsynaptic currents (uIPSCs) were evoked by inducing action potentials in RFP+ neurons. The amplitude and paired-pulse ratio (PPR), monitoring release probability, of uIPSCs and synaptic connectivity were compared between *Nlgn3Δ*-transfected and -untransfected neurons (**Figure 2B-E**). Importantly, VGT3+ inhibitory interneurons displayed clear potentiation of uIPSCs onto CA1 pyramidal neurons overexpressing Nlgn3Δ. Paired action potentials (APs) of VGT3+ neurons with short intervals (50 ms) induced paired-pulse depression (PPD) of uIPSCs. Nlgn3Δ displayed reduced PPD compared with untransfected neurons, consistent with previous work (Futai et al., 2007; Shipman et al., 2011; Uchigashima et al., 2020) (**Figure 2D**). As PPR inversely correlates with presynaptic release probability, these results suggest that Nlgn3Δ OE can facilitate presynaptic GABA release. In contrast, Nlgn3Δ OE reduced uIPSCs in Pv+ inhibitory synaptic transmission, but had no effect on PPR as reported previously (**Figure 3A-D**) (Horn & Nicoll, 2018). Lastly, Nlgn3Δ OE did not alter uIPSCs or PPR mediated by Sst+ interneurons (**Figure 3E-H**). No effect of biolistic transfection with EGFP alone was found on uIPSC amplitude, PPR or connection probability at Pv+ and Sst+ inhibitory synapses. The above results strongly suggest that Nlgn3 modifies inhibitory synaptic function depending on the type of presynaptic interneuron with which it interacts. Interestingly, Nlgn3Δ OE did not increase synaptic connectivity (**Figure 2E**), suggesting that i) Nlgn3 regulates pre-existing inhibitory inputs on postsynaptic neurons and/or ii) postsynaptic Nlgn3Δ OE is not sufficient to induce new synapse formation.

**Figure 2.**
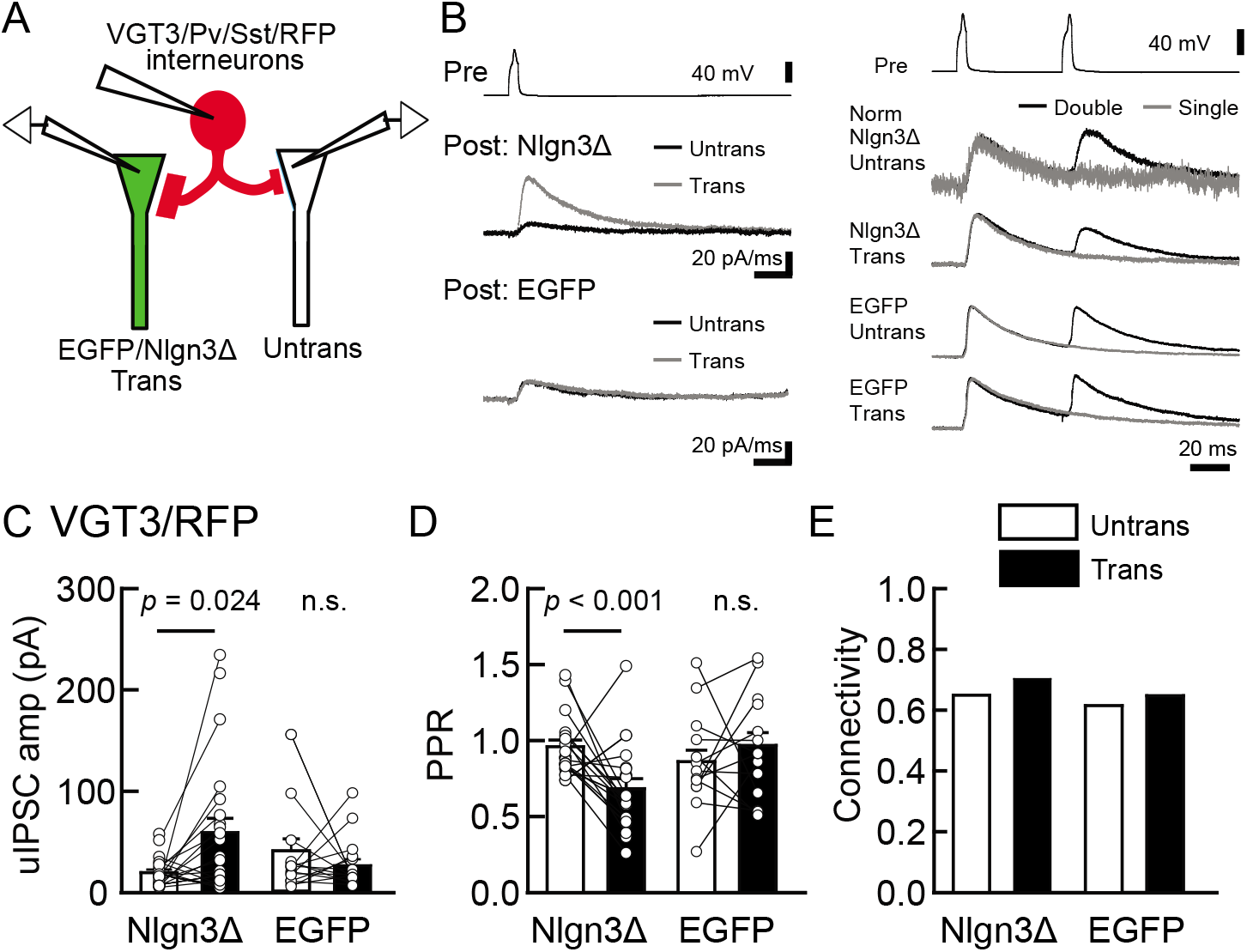
Nlgn3Δ overexpression specifically potentiates VGT3+ interneuron-mediated unitary synaptic transmission. The effects of overexpression of Nlgn3Δ isoform or EGFP (control) in hippocampal CA1 pyramidal neurons on inhibitory inputs mediated by VGT3+ inhibitory interneurons. **(A)** Configuration of the triple whole-cell recording for VGT3+, Pv+ and Sst+ interneurons forming synapses with pyramidal neurons. **(B)** Sample traces of uIPSCs. Left top, averaged sample traces of a single presynaptic action potential evoked in a VGT3+ interneuron. Left middle and bottom, superimposed averaged sample uIPSC traces (Untrans: black, trans: dark gray) induced by an action potential (AP). Right, superimposed averaged sample traces of uIPSCs evoked by single (dark gray) and double (black) APs in VGT3+ interneurons. uIPSCs are normalized to the first amplitude. Because the first uIPSC overlaps with the second uIPSC, to accurately measure the amplitude of the second IPSC, we ‘cancelled’ the first uIPSC by subtracting the traces receiving a single pulse (gray) from those receiving a paired pulse (black), both normalized to the first response. The amplitude **(C)** and paired-pulse ratio (PPR) **(D)** of uIPSCs were plotted for each pair of transfected (Trans) and neighboring untransfected (Untrans) cells (open symbols). Bar graphs indicate the mean ± SEM. **(E)** Synaptic connectivity between presynaptic inhibitory interneuron and postsynaptic untransfected (open bars) or transfected (black) pyramidal neurons. Numbers of cell pairs: Nlgn3Δ or EGFP at VGT3+ synapses (28 pairs/20 mice and 19/19). The number of tested slice cultures is the same as that of cell pairs. n.s., not significant. Mann–Whitney U test.

**Figure 3.**
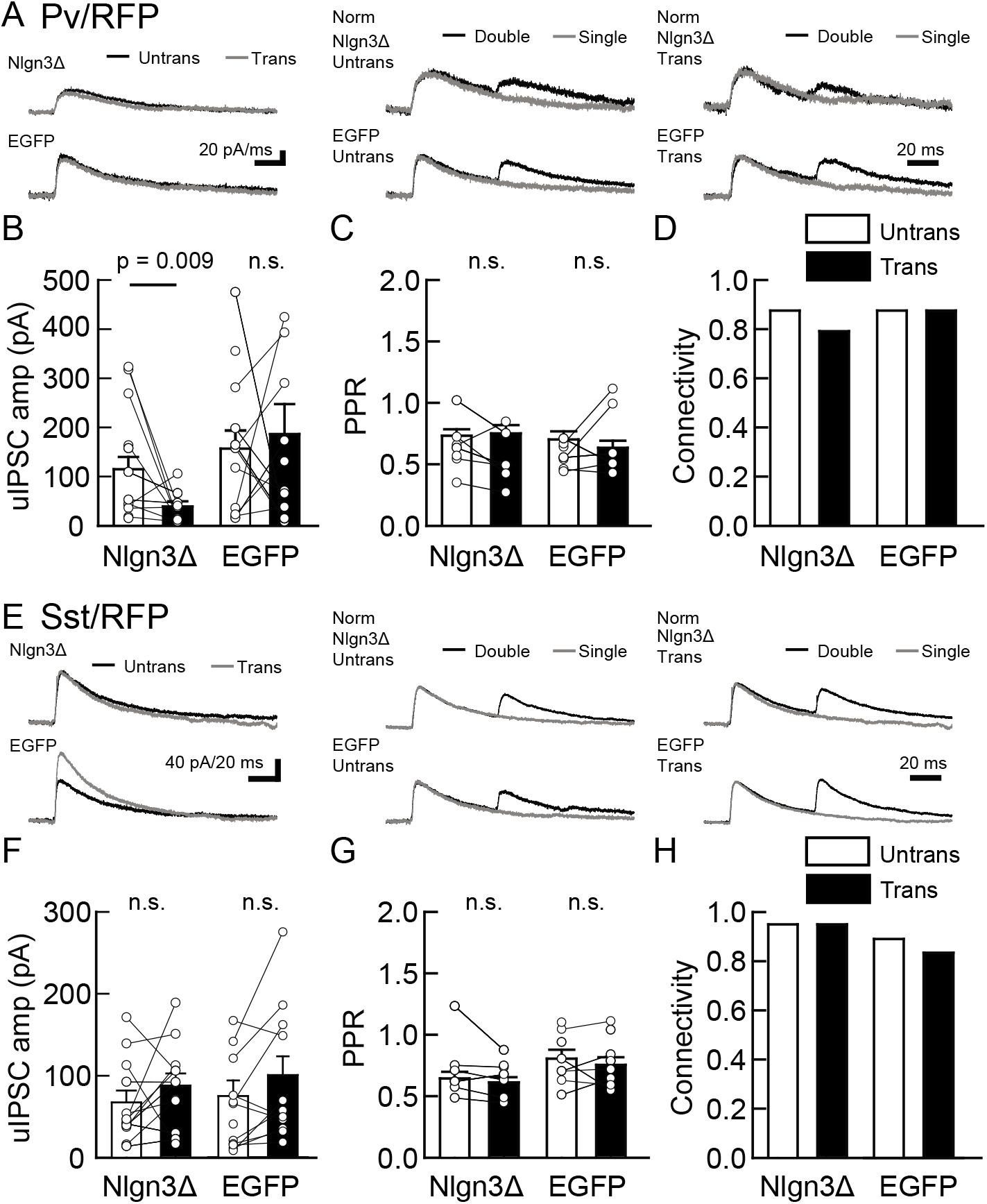
Nlgn3Δ overexpression does not increase PV+ and Sst+ interneuron-mediated unitary synaptic transmission. The effects of overexpression of Nlgn3Δ isoform or EGFP (control) in hippocampal CA1 pyramidal neurons were compared between inhibitory inputs mediated by Pv+ **(A-D)** and Sst+ **(E-H)** inhibitory interneurons. **(A and E)** Sample traces of uIPSCs. Left, superimposed averaged sample uIPSC traces (Untrans: black, trans: dark gray) induced by an AP. Middle and right, superimposed averaged sample traces of uIPSCs evoked by single (dark gray) and double (black) APs in Pv+ and Sst+ interneurons. uIPSCs are normalized to the first amplitude. The amplitude **(B and F)** and paired-pulse ratio (PPR) **(C and G)** of uIPSCs were plotted for each pair of transfected (Trans) and neighboring untransfected (Untrans) cells (open symbols). Bar graphs indicate the mean ± SEM. **(D and H)** Synaptic connectivity between presynaptic inhibitory interneuron and postsynaptic untransfected (open bars) or transfected (black) pyramidal neurons. Numbers of cell pairs: Nlgn3Δ or EGFP at Pv+ synapse (33 pairs/19 mice and 29/17) and Sst+ synapses (20/11 and 18/11). The number of tested slice cultures is the same as that of cell pairs. n.s., not significant. Mann–Whitney U test.

### Nlgn3 knockdown reduces VGT3+ inhibitory synaptic transmission in the hippocampal CA1 region

We next tested the impact of acute Nlgn3 knockdown (KD) on inhibitory synaptic transmission. Organotypic slice cultures prepared from C57BL/6J mice were biolistically transfected with shRNA against Nlgn3 (shNlgn3#1), which exhibits over 90% knockdown efficiency specific to Nlgn3 isoforms (**Figure S2A**), or control shRNA (shCntl). Transfection was performed at days *in vitro* (DIV) one and recordings were performed 7-9 days later to measure inhibitory synaptic transmission mediated by three different synaptic inputs (**Figure 4**). Compared with untransfected neurons, shNlgn3#1-transfected neurons displayed reduced uIPSC amplitudes mediated by VGT3+ interneurons (**Figure 4A-D**). In contrast, uIPSC amplitudes were affected in neither Pv+ (**Figure 4E-H**) nor Sst+ (**Figure 4I-L)** neurons. Another Nlgn3 KD shRNA, shNlgn3#2, also reduced uIPSC amplitudes at VGT3+ inhibitory synapses (**Figure S2B-F**), ruling out off-target effects of shRNAs. Neurons transfected with shCntl displayed uIPSC amplitudes comparable to untransfected neurons. No significant changes were detected in PPR (**Figure 4C, G, K**) and connection probability (**Figure 4D, H, L**) between transfected and untransfected neurons. Taken together, these data suggest that Nlgn3 is required for synaptic transmission specifically at VGT3+ GABAergic synapses in an input cell-dependent manner.

**Figure 4.**
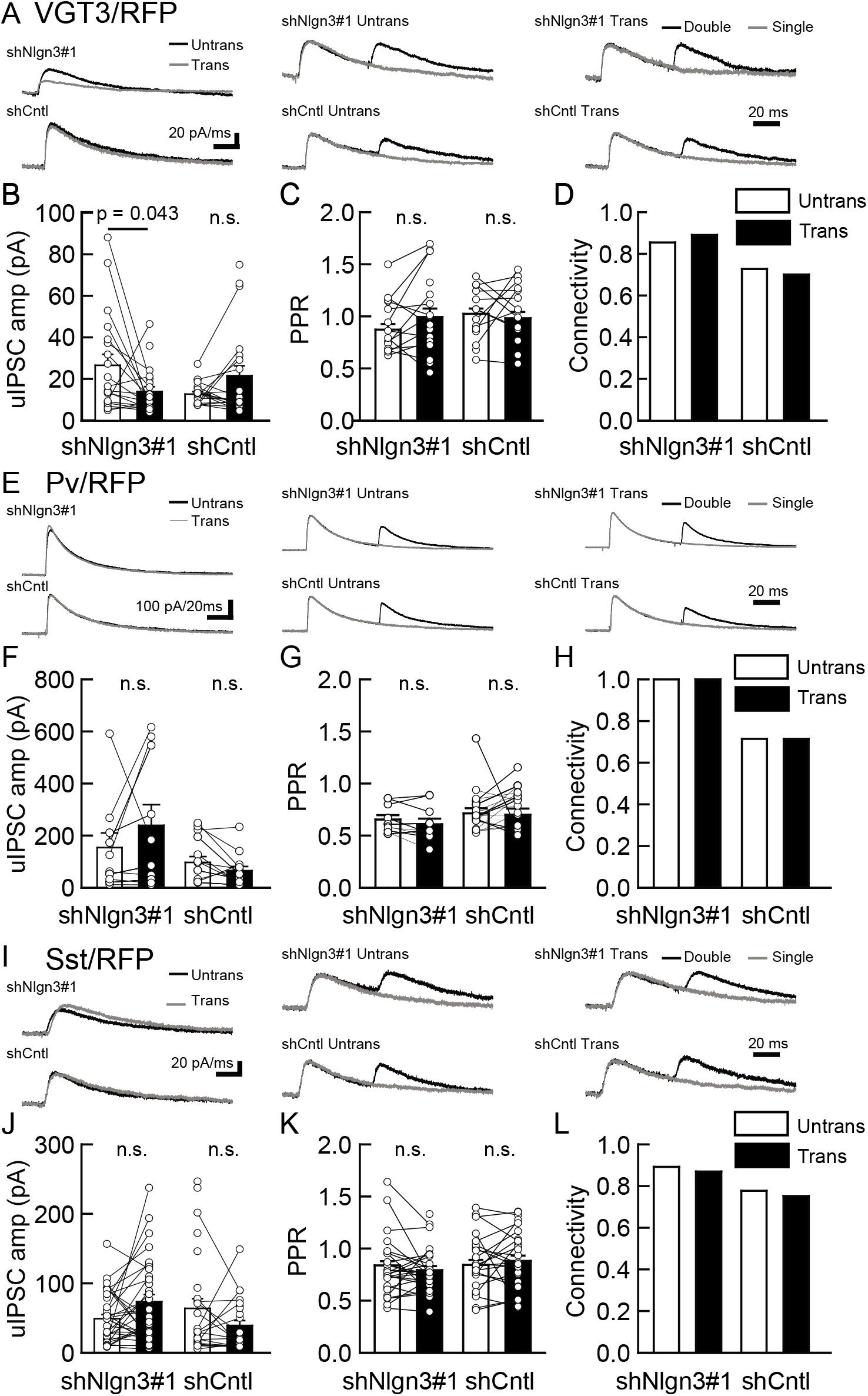
Endogenous Nlgn3 specifically regulates VGT3+, but not Pv+ and Sst+, interneuron-mediated unitary synaptic transmission. The effects of shNlgn3 or shCntl expression in hippocampal CA1 pyramidal neurons were compared between three different inhibitory inputs mediated by VGT3+ **(A-D)**, Pv+ **(E-H)** and Sst+ **(I-L)** inhibitory interneurons. **(A, E and I)** Sample traces of uIPSCs. Left, superimposed averaged sample traces of uIPSC (Untrans: black, trans: dark gray) induced by an AP. Middle and right, superimposed averaged sample traces of uIPSCs evoked by single (dark gray) and double (black) APs in VGT3+ (**A**), Pv+ (**E**) and Sst+ (**I**) interneurons. uIPSCs are normalized to the first amplitude. The amplitude **(B, F, J)** and PPR **(C, G, K)** of uIPSCs were plotted for each pair of transfected (Trans) and neighboring untransfected (Untrans) cells (open symbols). Bar graphs indicate the mean ± SEM. **(D, H, L)** Synaptic connectivity between presynaptic inhibitory interneuron and postsynaptic untransfected (open bars) or transfected (black) pyramidal neurons. Numbers of cell pairs: shNlgn3 or shCntl at VGT3+ synapses (28 pairs/15 mice and 37/17), at Pv+ synapse (12/4 and 16/5), and Sst+ synapses (26/11 and 21/10). The number of tested slice cultures is the same as that of cell pairs. n.s., not significant. Mann–Whitney U test.

### Lack of *Nrxn* genes in presynaptic VGT3+ neurons abolishes the effect of postsynaptic Nlgn3Δ OE

Our results suggest that Nlgn3 specifically translocates to VGT3+ inhibitory terminals and regulates inhibitory synaptic transmission. Postsynaptic Nlgns couple with presynaptic Nrxns to form trans-synaptic protein complexes that regulate synapse formation and function (Sudhof, 2017). Does input-specific Nlgn3Δ (**Figure 2C**) require presynaptic Nrxn proteins? To address this question, we generated VGT3 neuron-specific triple *Nrxn* knockout with TdTomato reporter gene mouse line (Nrxn1/2/3^f/f^/VGT3^cre^/TdTomato: NrxnTKO/VGT3/RFP). This mouse line is fertile with KO of *Nrxn1, 2* and *3* specifically in TdTomato-positive VGT3+ neurons (**Figure 5A**). We transfected Nlgn3Δ in NrxnTKO/VGT3/RFP slice cultures and performed triple whole-cell recordings as described above using VGT3/RFP slice cultures (**Figure 2B-E**). VGT3+ neurons lacking Nrxns induced synaptic release regardless of Nlgn3Δ gene transfection, indicating that presynaptic Nrxns in VGT3+ interneurons are not essential for synaptogenesis. Importantly, we observed no enhancement of uIPSC amplitude in Nlgn3-overexpressed neurons compared with untransfected pyramidal neurons (**Figure 5B-E**). These results strongly suggest that presynaptic Nrxn proteins are necessary for regulating inhibitory synaptic transmission through postsynaptic Nlgn3.

**Figure 5.**
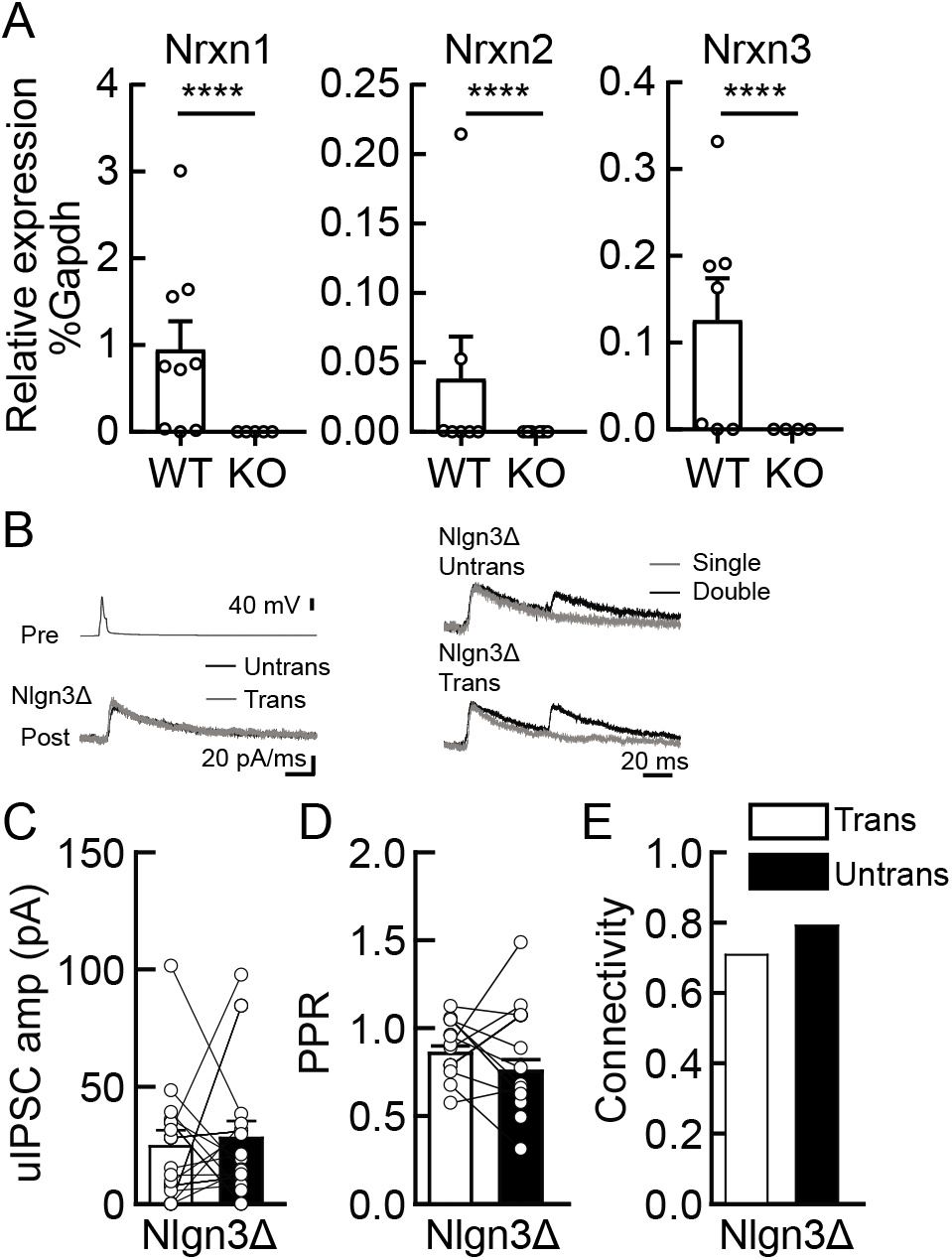
Lack of presynaptic Nrxns abolishes the potentiation effect of Nlgn3Δ in VGT3+ inhibitory synaptic transmission. Effects of Nlgn3Δ isoform overexpression in hippocampal CA1 pyramidal neurons in the absence of presynaptic Nrxn input in VGT3+ interneurons. **(A)** Validation of the NrxnTKO/VGT3/RFP mouse line. Expression of *Nrxn* genes were compared in TdTomato-positive VGT3+ neurons in organotypic hippocampal slice cultures prepared from WT (VGT3/RFP) and KO (NrxnTKO/VGT3/RFP) mice. qPCRs against *Nrxn 1, 2, 3* and *Gapdh* (internal control) were performed for single-cell cDNA libraries prepared from TdTomato-positive neurons. Number of neurons: WT (N = 9, 3 mice) and KO (7, 2). ****p < 0.0001 (Student’s *t*-test). **(B-E)** Effect of Nlgn3Δ OE on NrxnTKO VGT3+ interneuron-mediated inhibitory synaptic transmission. **(B)** Left top, averaged sample traces of a single presynaptic AP evoked in a NrxnTKO VGT3+ interneuron. Left bottom, superimposed averaged sample traces of uIPSC (Untrans: black, trans: dark gray) induced by an AP. Right, superimposed averaged sample traces of uIPSCs evoked by single (dark gray) and double (black) APs in NrxnTKO VGT3+ interneurons. Summary of uIPSC amplitude **(C)**, PPR **(D)** and connectivity **(E)**. Open circles connected with bars represent individual pairs of cells **(C, D)**. Bar graphs indicate the mean ± SEM. N = 18 cell pairs (3 mice). The number of tested slice cultures is the same as that of cell pairs. Mann–Whitney U test.

### *αNrxn1* and *βNrxn3* mRNAs are highly expressed in VGT3+ interneurons

Our results above clearly suggest that presynaptic Nrxn proteins are important for the function of postsynaptic Nlgn3Δ. We therefore hypothesized that *Nrxn* isoforms highly expressed in VGT3+ interneurons functionally couple with postsynaptic Nlgn3Δ. To address this hypothesis, we examined the mRNA expression patterns of α and β isoforms of *Nrxn1–3* in hippocampal CA1 interneurons by fluorescent *in situ* hybridization (FISH). The specificities of cRNA probes for 6 *Nrxn* isoforms were validated recently (Uchigashima, Cheung, Suh, Watanabe, & Futai, 2019). mRNAs encoding all *Nrxn* isoforms, except *βNrxn1*, were detected not only in the st. pyramidale but also in scattered cells across all layers (**Figure S3A-F**). Importantly, *αNrxn1* and *βNrxn3* mRNAs appeared to be enriched in scattered cells within the st. radiatum or pyramidale (arrows in **Figure 6A-C and G-I**), where VGT3+ interneurons are dominantly distributed compared with Pv+ and Sst+ interneurons (Pelkey et al., 2017). Double FISH signals were twice as strong for *αNrxn1* and *βNrxn3* mRNAs in VGT3+ (**Figure 6A, G, J**) interneurons than in Pv+ (**Figure 6B, H, J**) and Sst+ (**Figure 6C, I and J**) interneurons. In contrast, there were no differences in the signal intensities for the remaining *Nrxn* isoforms between VGT3+ and other interneurons (**Figure 6D-F and J, S3**). These findings suggest that *αNrxn1* and *βNrxn3* mRNAs are highly expressed in VGT3+ interneurons compared with Pv+ or Sst+ interneurons.

**Figure 6.**
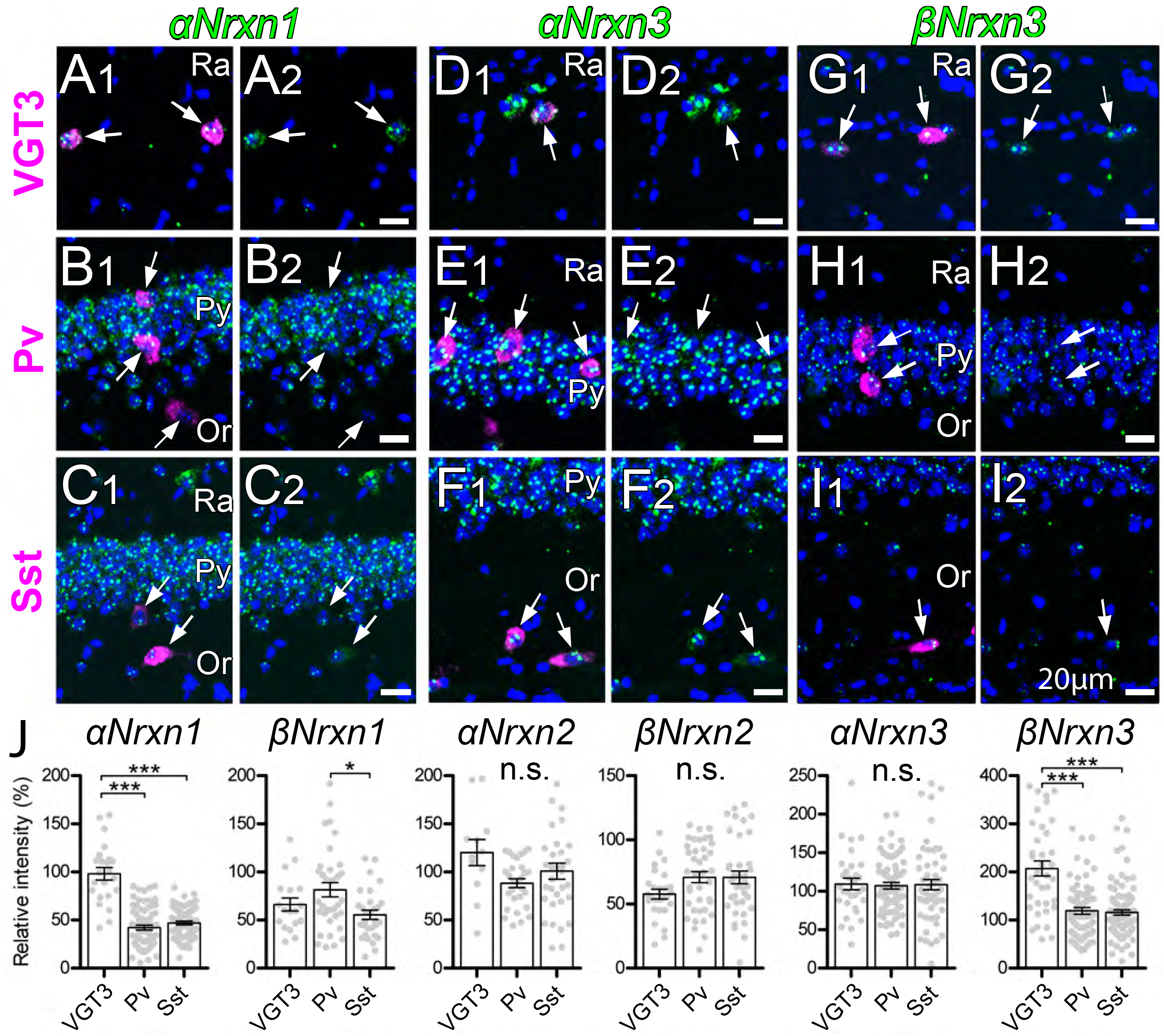
Expression of *Nrxn* isoforms in different types of inhibitory interneurons in the hippocampal CA1 region. Double FISH for αNrxn1 **(A, B, C)**, αNrxn3 **(D, E, F)** and βNrxn3 **(G, H, I)** in VGT3+ **(A, D, G)**, Pv+ **(B, E, H)** and Sst+ **(C, F, I)** in the hippocampus showing different levels of Nrxn mRNA (green) expression in different inhibitory interneurons (magenta). Note that the signal intensity in individual GABAergic neurons is variable, compared with that in glutamatergic neurons. Nuclei were stained with DAPI (blue). Or, st. oriens; Py, pyramidal cell layer; Ra, st. radiatum. **(J)** Summary scatter plots for six Nrxn mRNA in VGT3+, Pv+ and Sst+ inhibitory interneurons. Data are represented as mean ± SEM. ns, not significant; *p < 0.05; ***p < 0.001 (Mann–Whitney U test). Scale bars, 20 μm.

### Expression profiles of *Nrxn* splice isoforms in hippocampal VGT3+ inhibitory interneurons

Single-cell RNA sequencing was performed to elucidate splice variant expression of *Nrxn* isoforms in VGT3+ interneurons. We harvested cytosol from four VGT3/RFP neurons through whole-cell glass electrodes and performed single-cell deep RNA-seq. The t-SNE plot indicates that the genome-wide transcriptomes of the four cells (denoted as G556) were clustered together and close to that of adult GABAergic neurons derived from the single-cell RNA-seq dataset retrieved from the Allen Brain Map Cell Types Database (Mouse - Cortex and Hippocampus dataset, **Figure 7**). The expression of *Nrxn* genes in these four cells was similar to that of GABAergic neurons (**Figure 7A**). The quantification of *Nrxn* genes indicates that the expression of *Nrxn1* and *Nrxn3* are dominant in VGT3+ interneurons (**Figure 7 C and S4**). We then compared the expression of *Nrxn* splice isoforms in each *Nrxn* gene. Given that the insertion of AS4 determines the binding of many Nrxn protein binding partners including Nlgns (Sudhof, 2008, 2017), we quantified the expression of *Nrxn* isoforms with or without AS4 insertion. Twelve *Nrxn* splice isoforms, *αNrxn1+AS4, αNrxn1-AS4, αNrxn2+AS4, αNrxn2-AS4, αNrxn3+AS4, αNrxn3-AS4, βNrxn1+AS4, βNrxn1-AS4, βNrxn2+AS4, βNrxn2-AS4, βNrxn3+AS4* and *βNrxn3-AS4* were manually modified (**Figure 7C-E, Table 1**), and their expression was compared. Among *Nrxn1* splice isoforms, *αNrxn1+AS4* was consistently detected as the sole *αNrxn1* gene expressed in VGT3+ interneurons (**Figure 7C**). *βNrxn2+AS4* was the only confirmed *Nrxn2* gene expressed in VGT3+ neurons but its expression was much lower than that of *Nrxn1* and *3* (**Figure 7D**). *αNrxn3+AS4* and *βNrxn3-AS4* were the two major *Nrxn3* genes expressed in VGT3+ interneurons (**Figure 7E**). Our two gene expression assays, double FISH and single-cell RNA-seq, suggest that *αNrxn1* and *βNrxn3*, but not *αNrxn3*, are the major *Nrxn* genes expressed in VGT3+ neurons compared with PV+ and Sst+ neurons (**Figure 6J**), and *αNrxn1+AS4, αNrxn3+AS4* and *βNrxn3-AS4* are *Nrxn* splice isoforms highly expressed in VGT3+ neurons (**Figure 7C and E**). Therefore, *αNrxn1+AS4* and *βNrxn3-AS4* are the unique *Nrxn* genes expressed in VGT3+ neurons compared with PV+ and Sst+ neurons.

**Figure 7.**
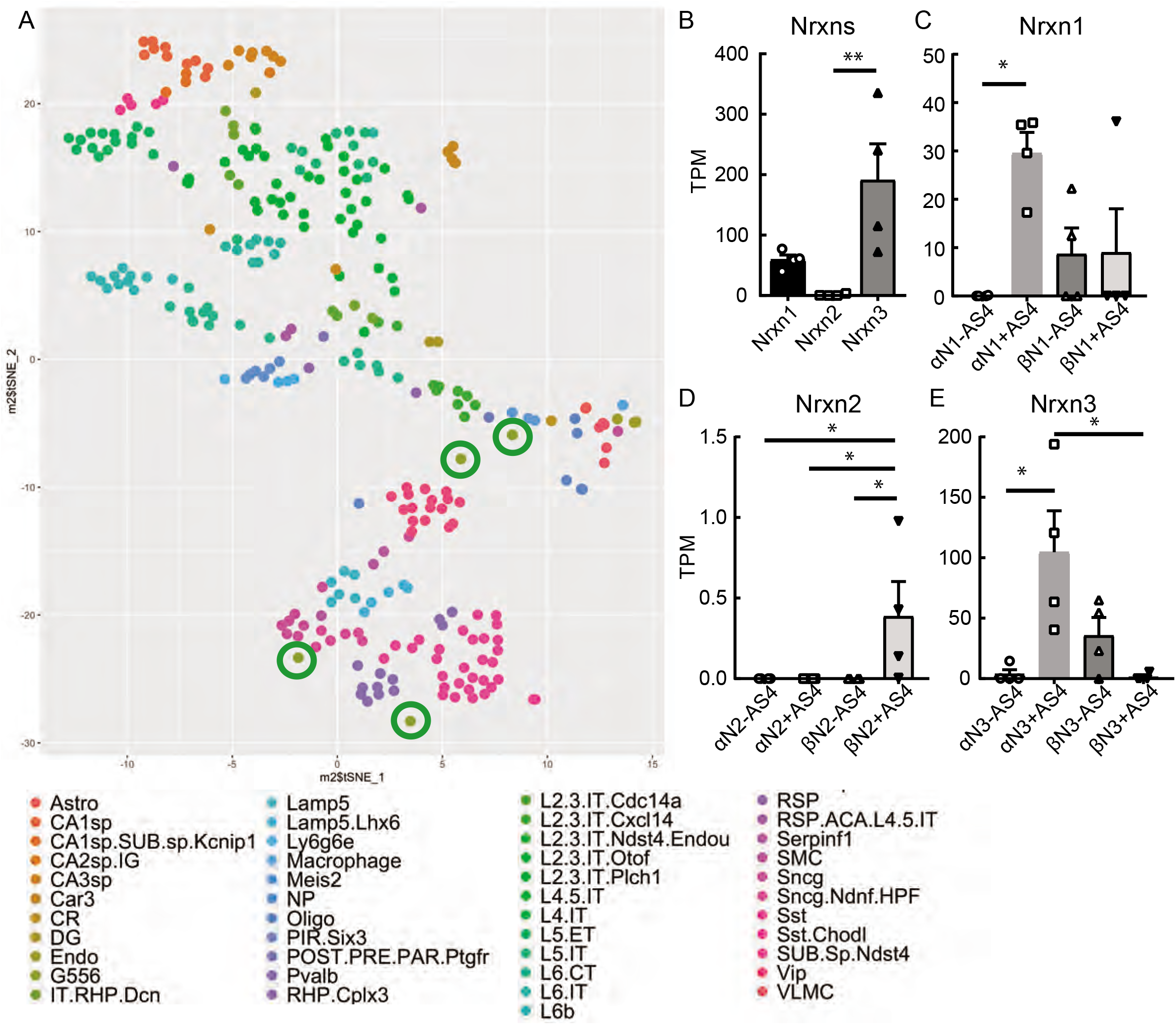
Endogenous *Nrxn* expression in hippocampal VGT3+ inhibitory interneurons. **(A)** Single-cell t-SNE plot of four single-cells (G556.06, .07, .09 and .10, green circles) compared to Allen Brain Atlas single-cells. **(B)** Summary bar graph of *Nrxn* gene (*Nrxn1, 2* and *3*) expression. **(C-E)** Summary graphs of splice isoforms of *Nrxn1* **(C)**, *2* **(D)** and *3* **(E)**. *p < 0.05, one-way ANOVA followed by Sidak’s multiple comparisons test or U-test, N = 4.

**Table 1.**
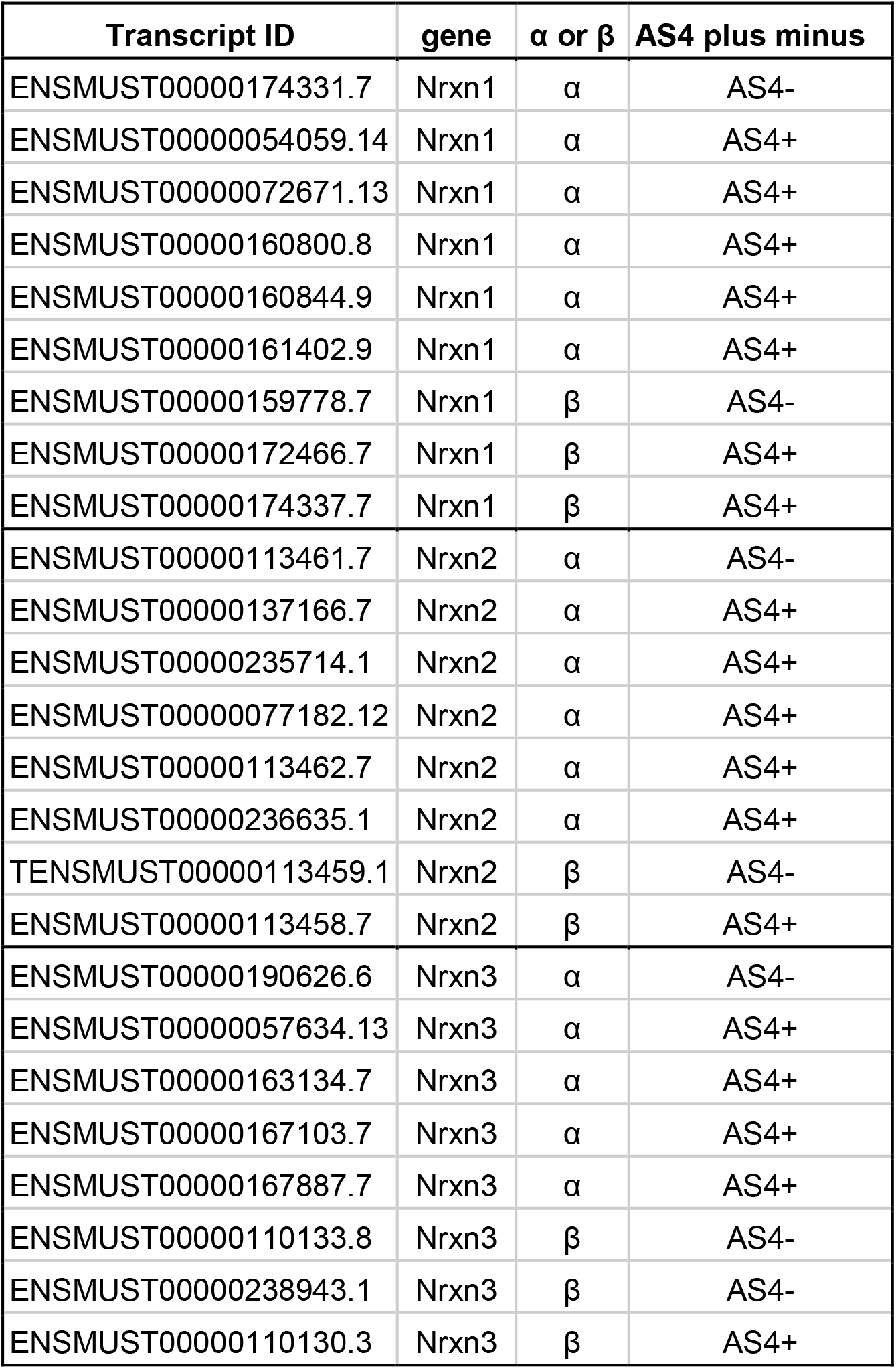
Nrxn transcript IDs used for quantification.

| **Transcript ID** | **gene** | **$\alpha$ or $\beta$** | **AS4 plus minus** |
| --- | --- | --- | --- |
| ENSMUST00000174331.7 | Nrxn1 | $\alpha$ | AS4- |
| ENSMUST00000054059.14 | Nrxn1 | $\alpha$ | AS4+ |
| ENSMUST00000072671.13 | Nrxn1 | $\alpha$ | AS4+ |
| ENSMUST00000160800.8 | Nrxn1 | $\alpha$ | AS4+ |
| ENSMUST00000160844.9 | Nrxn1 | $\alpha$ | AS4+ |
| ENSMUST00000161402.9 | Nrxn1 | $\alpha$ | AS4+ |
| ENSMUST00000159778.7 | Nrxn1 | $\beta$ | AS4- |
| ENSMUST00000172466.7 | Nrxn1 | $\beta$ | AS4+ |
| ENSMUST00000174337.7 | Nrxn1 | $\beta$ | AS4+ |
| ENSMUST00000113461.7 | Nrxn2 | $\alpha$ | AS4- |
| ENSMUST00000137166.7 | Nrxn2 | $\alpha$ | AS4+ |
| ENSMUST00000235714.1 | Nrxn2 | $\alpha$ | AS4+ |
| ENSMUST00000077182.12 | Nrxn2 | $\alpha$ | AS4+ |
| ENSMUST00000113462.7 | Nrxn2 | $\alpha$ | AS4+ |
| ENSMUST00000236635.1 | Nrxn2 | $\alpha$ | AS4+ |
| TENSMUST00000113459.1 | Nrxn2 | $\beta$ | AS4- |
| ENSMUST00000113458.7 | Nrxn2 | $\beta$ | AS4+ |
| ENSMUST00000190626.6 | Nrxn3 | $\alpha$ | AS4- |
| ENSMUST00000057634.13 | Nrxn3 | $\alpha$ | AS4+ |
| ENSMUST00000163134.7 | Nrxn3 | $\alpha$ | AS4+ |
| ENSMUST00000167103.7 | Nrxn3 | $\alpha$ | AS4+ |
| ENSMUST00000167887.7 | Nrxn3 | $\alpha$ | AS4+ |
| ENSMUST00000110133.8 | Nrxn3 | $\beta$ | AS4- |
| ENSMUST00000238943.1 | Nrxn3 | $\beta$ | AS4- |
| ENSMUST00000110130.3 | Nrxn3 | $\beta$ | AS4+ |

### Presynaptic αNrxn1+AS4 couples with postsynaptic Nlgn3Δ to regulate inhibitory function

Our electrophysiology results using NrxnTKO/VGT3/RFP mice indicate that Nrxn proteins are important for postsynaptic Nlgn3Δ function. We sought to determine which Nrxn isoform(s) is the functional partner(s) of Nlgn3Δ at VGT3+ inhibitory synapses. Does VGT3+ interneuron-dominant *Nrxns, αNrxn1* and *βNrxn3*, interact with postsynaptic Nlgn3Δ to modulate inhibitory synaptic function? To address this, we performed a rescue approach by expressing specific *Nrxn* isoforms in NrxnTKO/VGT3/RFP neurons. We expressed *Nrxn* with tagBFP and *Nlgn3Δ* with EGFP in VGT3/RFP and pyramidal neurons, respectively, in NrxnTKO/VGT3/RFP slice cultures using our recently developed electroporation technique (Keener, Cheung, & Futai, 2020), and performed triple whole-cell recordings from untransfected and tagBFP/Nrxn-transfected VGT3/RFP neurons, and GFP/*Nlgn3Δ*-transfected pyramidal neurons (**Figure 8**). We first identified two neighboring VGT3/RFP neurons in the hippocampal st. radiatum and oriens regions and electroporated tagBFP/Nrxn plasmids into one of the cells, while *GFP/Nlgn3Δ* was electroporated into pyramidal neurons near the electroporated VGT3/RFP neuron. Given that splice insertion at site 4 in Nrxns regulates binding with postsynaptic Nlgns (Sudhof, 2008, 2017), we transfected four *Nrxn* splice isoforms, *αNrxn1+AS4, αNrxn1-AS4, βNrxn3+AS4 or βNrxn3-AS4*, together with tagBFP into VGT3/RFP neurons in CA1 st. radiatum and pyramidale. Two to three days after electroporation, we performed triple whole-cell recording from *Nrxn*-transfected presynaptic interneurons (Tdtomato- and tagBFP-positive) located in close proximity to untransfected VGT3/RFP (**Figure 8A**) and *Nlgn3Δ*-transfected (EGFP-positive) postsynaptic pyramidal neurons. Cell pairs overexpressing *αNrxn1+AS4* and *Nlgn3Δ* in presynaptic VGT3/RFP and postsynaptic pyramidal neurons, respectively, displayed significant enhancement of uIPSC with 100% connectivity (**Figure 8B, C, E**). In contrast, presynaptic *αNrxn1-AS4/tagBFP* and postsynaptic GFP/Nlgn*3Δ* neuron pairs showed no enhancement of uIPSC compared with control cell pairs, suggesting that Nlgn3Δ specifically couples with αNrxn1+AS4 to regulate inhibitory synaptic function at VGT3+ synapses. Transfection of tagBFP into VGT3/RFP neurons did not alter inhibitory synaptic transmission. Next, we tested βNrxn3-Nlgn3Δ synaptic codes on VGT3+ inhibitory synaptic transmission (**Figure 9A-D**). In contrast with αNrxn1-Nlgn3Δ codes, neither *βNrxn3+AS4* nor *βNrxn3-AS4* transfection showed detectable changes in uIPSCs compared with untransfected VGT3/RFP neurons, indicating that βNrxn3 is not important for Nlgn3-mediated inhibitory synaptic function. Interestingly, βNrxn3-AS4 expressed in NrxnTKO/VGT3/RFP/tagBFP neurons demonstrated reduced connectivity (**Figure 9D**), suggesting that βNrxn3-AS4 protein hinders synapse formation between VGT3/RFP terminals and postsynaptic CA1 pyramidal neurons.

**Figure 8.**
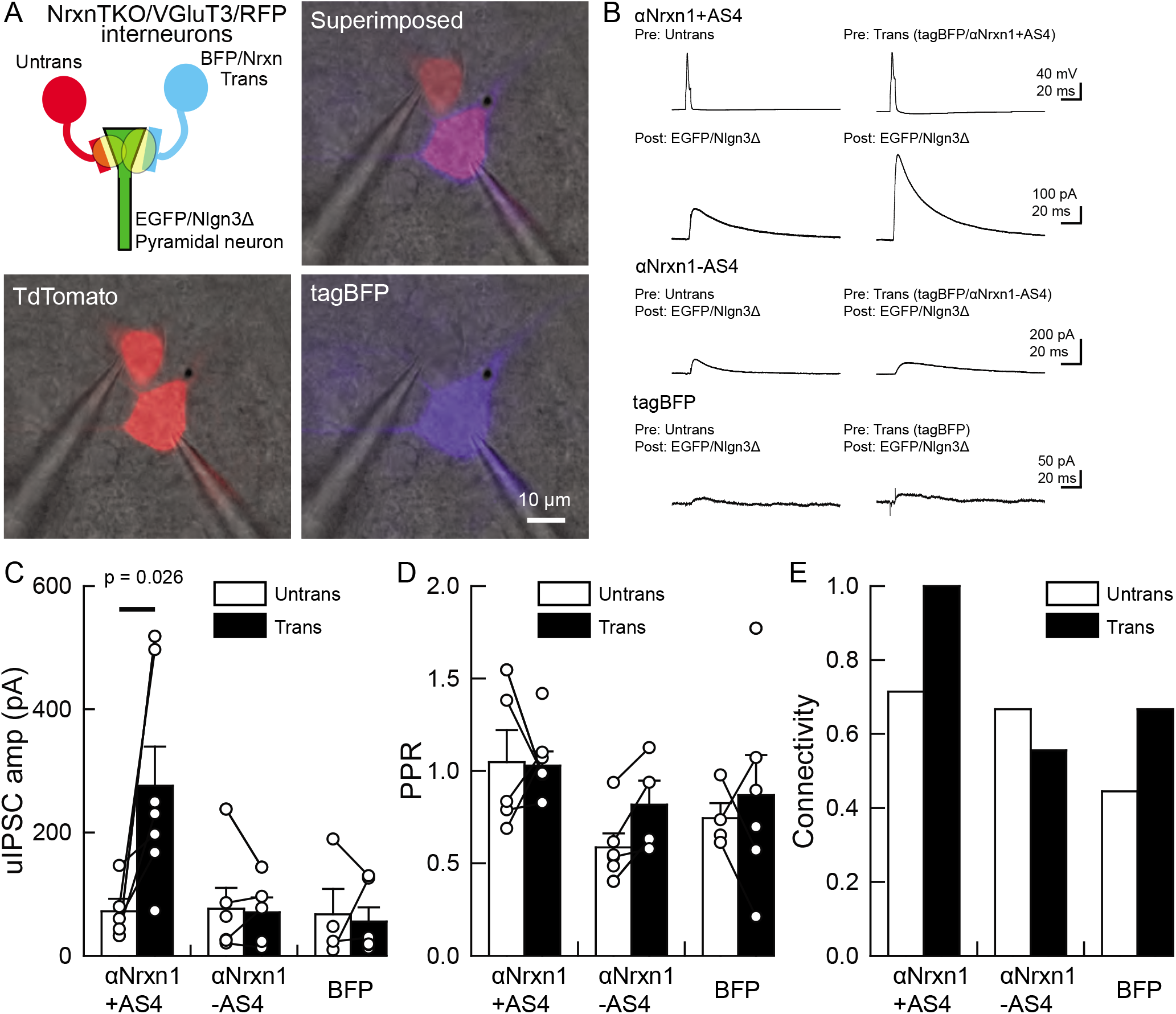
An αNrxn1+AS4–Nlgn3Δ synaptic code enhances VGT3+ inhibitory synapse transmission. **(A–C)** Effect of pre- and postsynaptic overexpression of αNrxn1 and Nlgn3Δ, respectively, on unitary inhibitory synaptic transmission in organotypic slice cultures prepared from VGT3/NrxnTKO/RFP mice. **(A)** Configuration of triple whole-cell recording (left top), superimposed fluorescent and Nomarski images (right top). (Bottom) Individual fluorescence and Nomarski images of TdTomato (left) and tagBFP (right). **(B)** Averaged sample uIPSC traces. Nrxns and Nlgn3Δ were transfected in TdTomato-positive and CA1 pyramidal neurons, respectively, by electroporation. **(C, D, E)** Summary of uIPSC amplitude **(C)**, paired-pulse ratio **(D)** and connectivity **(E)**. Numbers of cell pairs: αNrxn1+AS4, αNrxn1-AS4, and BFP at VGT3+ synapses (7 pairs/3 mice, 9/4, and 9/4). The number of tested slice cultures is the same as that of cell pairs. Mann–Whitney U test.

**Figure 9.**
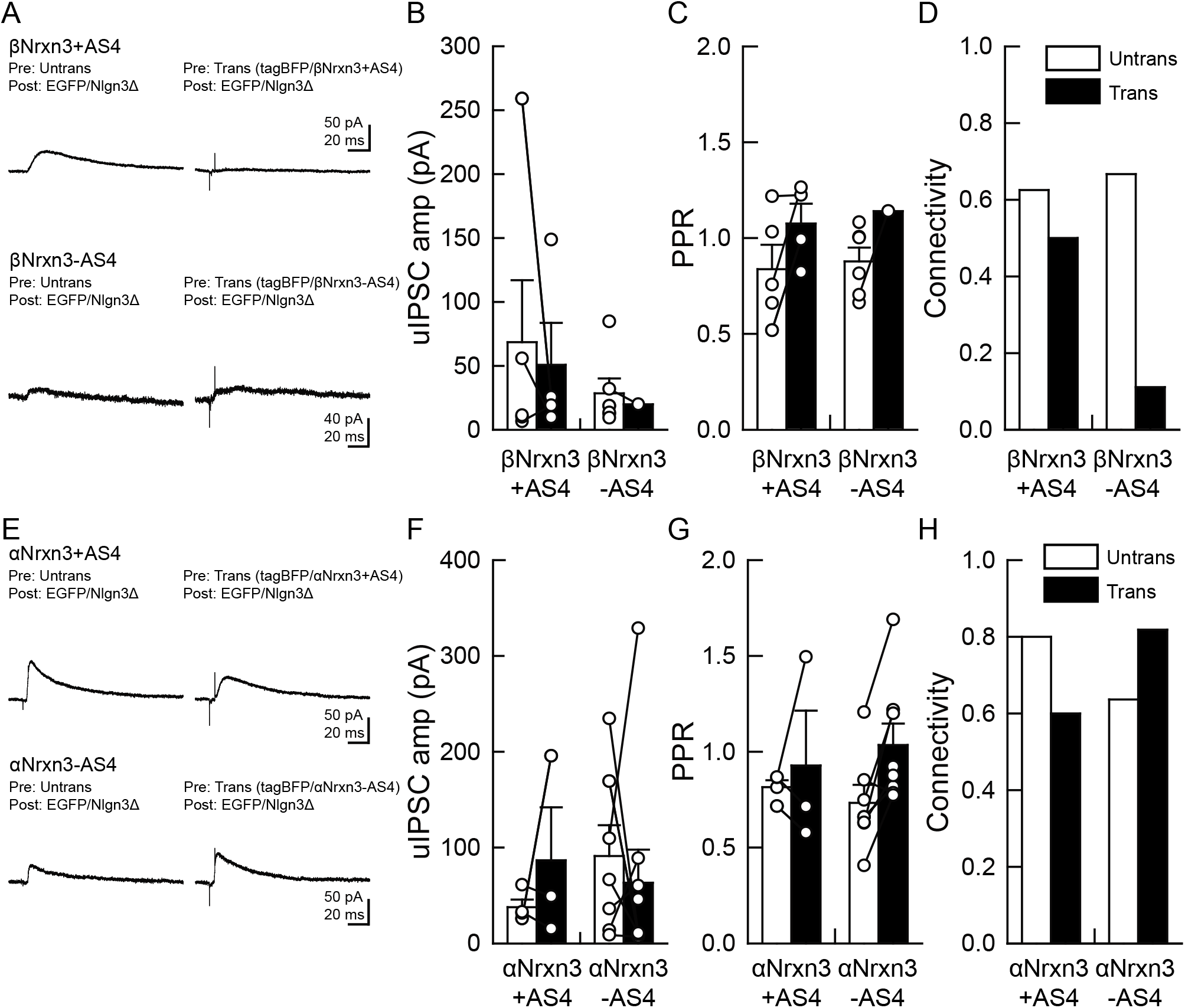
α/βNrxn3–Nlgn3Δ synaptic codes do not regulate VGT3+ inhibitory synapse transmission. **(A-D)** Effect of pre- and postsynaptic overexpression of βNrxn3(±AS4) and Nlgn3Δ, respectively, on unitary inhibitory synaptic transmission in organotypic slice cultures prepared from NrxnTKO/VGT3/RFP mice. **(E-H)** Effect of pre- and postsynaptic overexpression of αNrxn3(±AS4) and Nlgn3Δ, respectively, on unitary inhibitory synaptic transmission. **(A, E)** Averaged sample uIPSC traces. Summary of uIPSC amplitude **(B, F)**, paired-pulse ratio **(C, G)** and connectivity **(D, H)**. Numbers of cell pairs: βNrxn3+AS4, βNrxn3-AS4, αNrxn3+AS4, and αNrxn3-AS4 at VGT3+ synapses (8 pairs/3 mice, 9/3, 5/3, and 11/3). The number of tested slice cultures is the same as that of cell pairs. Mann–Whitney U test.

Our rescue approach suggests that αNrxn+AS4 regulates inhibitory synaptic transmission with Nlgn3Δ. The gene structures of α and βNrxns are similar, therefore, different *αNrxn+AS4* isoform(s) may be able to functionally substitute αNrxn1+AS4. Our FISH results indicate that VGT3+, PV+ and Sst+ interneurons express comparable levels of *αNrxn3* (**Figure 6D-F and J**). Thus, we tested whether *αNrxn3+/-AS4* functionally couples with *Nlgn3Δ* in NrxnTKO/VGT3/RFP slice cultures (**Figure 9E-H**). To our surprise, neither *αNrxn3+AS4* nor *αNrxn3-AS4* pairing with postsynaptic Nlgn3Δ had any effect on inhibitory synaptic transmission. These results strongly suggest that αNrxn1+AS4, but not αNrxn3+AS4, has a unique code in the extracellular domain important for synaptic function with Nlgn3Δ.

## Discussion

Synaptic protein-protein interactions are critical for the development, maturation and survival of neurons. However, it is technically challenging to physiologically characterize trans-synaptic CAM protein interactions in two different neurons due to the difficulty in identifying synaptically-connected neuronal pairs in the brain. Co-culture approaches consisting of non-neuronal cells transfected with different *Nrxn* splice isoforms and dissociated neurons expressing endogenous Nlgns (or expressing Nrxn-binding proteins in non-neuronal cells and observing their interactions with endogenous Nrxns) have begun to elucidate the roles of trans-synaptic Nrxn/Nlgn isoforms on the clustering of pre- /postsynaptic molecules (Chih et al., 2006; Kang, Zhang, Dobie, Wu, & Craig, 2008; Ko et al., 2009; Nam & Chen, 2005; Scheiffele, Fan, Choih, Fetter, & Serafini, 2000). However, this approach is limited to primary cultures and cannot address whether these trans-synaptic interactions are sufficient to induce functional synapse diversification. To fully understand the physiological roles of trans-synaptic molecules, one must be able to manipulate the expression of these molecules in pre- and postsynaptic neurons simultaneously followed by determination of the functional consequences of such manipulation. Using our newly developed gene electroporation method that enables us to transfect genes in minor cell types such as specific inhibitory interneurons (Keener et al., 2020), we demonstrated for the first time that αNrxn1+AS4 and Nlgn3Δ, which are endogenously expressed in VGT3+ inhibitory and CA1 pyramidal neurons (**Figure 7D**) (Futai et al., 2007; Shipman et al., 2011; Uchigashima et al., 2020), respectively, form a specific code that dictates inhibitory synaptic transmission.

It has been reported that Nlgn proteins regulate inhibitory synaptic transmission in an input cell-specific manner. For example, a Nlgn2 KO mouse line displays deficits in fast-spiking but not Sst+ interneuron-mediated inhibitory synaptic transmission in the somatosensory cortex (Gibson, Huber, & Sudhof, 2009). Furthermore, Nlgn3 KO mice and the Nlgn3 R451C knock-in mutant line, which mimics a human autism mutation, showed Pv+ or Cck+ input-specific abnormal inhibitory synaptic transmission in the hippocampal CA1 region (Foldy et al., 2013). Therefore, the function of postsynaptic Nlgns are determined by the type of presynaptic inputs it receives, supporting the intriguing hypothesis that specific Nrxn-Nlgn binding regulates synaptic function. Our gene expression and functional assays highlight that *αNrxn1* is abundantly expressed in VGT3+ interneurons compared with other interneurons (**Figure 6J**) and *αNrxn1+AS4* is the dominant *αNrxn1* splice isoform expressed in VGT3+ interneurons (**Figure 7C**) which regulates inhibitory synaptic transmission with Nlgn3Δ (**Figure 8C**). On the other hand, our FISH experiments, which do not distinguish the AS4 insertion, detected *αNrxn1* at low levels in Pv+ and Sst+ interneurons (**Figure 6J**). The expression of *αNrxn1+AS4* was also previously confirmed in Pv+ interneurons (Fuccillo et al., 2015). Considering the lower expression of Nlgn3 at Pv+ inhibitory synapses (**Figure 1L**), other postsynaptic mechanism(s) might exist to regulate inhibitory synaptic transmission with αNrxn1+AS4 at Pv+ inhibitory synapses.

Biochemical studies have demonstrated that any Nrxn can bind to any Nlgn with different affinities, with one exception, αNrxn1-Nlgn1 interaction (Boucard et al., 2005; Reissner et al., 2008). However, our results clearly showed that αNrxn1+AS4, but not the other Nrxn variants we analyzed, contributes to Nlgn3Δ-mediated function at VGT3+ synapses. Similarly, αNrxn1-AS4 is known to specifically interact with Nlgn2 and enhances inhibitory synaptic transmission (Futai et al., 2013). This gap between biochemical binding affinities and physiological synaptic functions is likely due to interactions of Nrxn and Nlgn with other synaptic proteins. In this regard, our study dissects physiological Nrxn-Nlgn interactions at synapses compared with biochemical analyses of purified Nrxn and Nlgn proteins.

The AS4 is the most important splice insertion site in Nrxns to change the affinity to postsynaptic binding partners, including Nlgn (Sudhof, 2017). A splice insertion at AS4 of βNrxn1 can weaken the interaction with Nlgns (Koehnke et al., 2010). Indeed, we have reported that βNrxn1-AS4 but not βNrxn1+AS4 increases synaptic transmission via interaction with Nlgn1 (Futai et al., 2013). In contrast, we found that an insertion in AS4 of αNrxn1 can increase inhibitory synaptic transmission at VGT3+ synapses via trans-synaptic interaction with Nlgn3Δ. It is particularly interesting that only αNrxn1+AS4, but not αNrxn3+AS4, encoded uIPSC enhancement with postsynaptic Nlgn3Δ (**Figure 8C and 9F**). These results suggest that structural differences beyond the AS4 site exist between these two αNrxns. Since the structures of the major Nrxn domains, such as LNS and EGFA-EGFC, are similar between αNrxns, one possible interpretation is that differential alternative splicing events occurring at other AS sites may regulate binding with Nlgn3Δ. Further structural and functional analyses targeting these AS events could reveal a novel AS structure that dictates Nrxn-Nlgn codes on synaptic function. Moreover, it will be important to address whether αNrxn1+AS1, αNrxn3+AS4 and βNrxn3-AS4, which are highly expressed in VGT3+ neurons (**Figure 7**), can encode specific synaptic functions when interacting with other postsynaptic Nrxn-binding partners such as Nlgn2, CST-3 and IgSF21 (Pettem et al., 2013; Tanabe et al., 2017; Um et al., 2014).

Deletion of Nrxns or Nlgns has been reported to have little effect on synapse formation (Chanda, Hale, Zhang, Wernig, & Sudhof, 2017; Chen, Jiang, Zhang, Gokce, & Sudhof, 2017; Varoqueaux et al., 2006). Our manipulation of the expression levels of either presynaptic Nrxns or postsynaptic Nlgn3 consistently did not affect synaptic connectivity, a measurement of the number of active synapses (**Figure 2, 3 and 6**). However, Nlgn3Δ formed new synapses only when αNrxn1+AS4 was simultaneously expressed in VGT3+ interneurons (**Figure 8E**). This suggests that trans-synaptic interactions of Nrxns and Nlgns may control not only synapse function but also synapse number, while the lack of Nrxns or Nlgns on either side of the synapse can be compensated by other CAM interactions. In this context, it is important to address whether the subcellular localization of Nrxn and Nlgn proteins is regulated by synaptic activity, such as homeostatic synaptic plasticity (Mao et al., 2018).

Note that our results, demonstrating the role of postsynaptic Nlgn3Δ in CA1 pyramidal neurons on inhibitory synaptic transmission, have some inconsistencies with a previously published study, which found that Nlgn3 differentially regulates Pv+ and Sst+ interneuron inhibitory synapses (Horn & Nicoll, 2018). Horn and Nicoll reported that OE of human Nlgn3A2 reduced Pv+ and increased Sst+ inhibitory synaptic transmission. The former finding is consistent with our results in **Figure 3B**, displaying that Nlgn3Δ OE reduces Pv+ uIPSC amplitude. This may suggest that Pv+ neurons have a unique trans-synaptic regulatory mechanism compared with other interneurons and that Nlgn3Δ OE may disrupt endogenous GABAAR complexes at Pv+ inhibitory synapses. In contrast, Nlgn3Δ OE did not increase Sst+ uIPSCs (**Figure 3F**), as observed in Horn & Nicoll, 2018. This difference might be due to variations in experimental approaches including the Nlgn3 clone used (human Nlgn3A2 versus mouse Nlgn3Δ slice isoform) and the duration of transgene or shRNA expression in hippocampal CA1 pyramidal neurons. Additionally, Horn and Nicoll crossed Pv^cre^ and Sst^cre^, the same *cre* lines we tested, with Ai32 mice, a *cre*-dependent channelrhodopsin line (Madisen et al., 2012), to evoke Pv+ or Sst+ neuron-mediated synaptic transmission, respectively (Horn & Nicoll, 2018). However, our current and previous studies for mouse line validation indicate that while PV^cre^ and Sst^cre^ lines exhibit highly specific *cre* expression in these cell types, these lines have leaky *cre* expression in other cell type(s) (**Figure S1**, yellow arrow heads) (Mao et al., 2015). Therefore, light-evoked activation of non-specific cell types may contribute to the inconsistent results in synaptic transmission.

Mutations and deletions in Nrxn loci are associated with neuropsychiatric and neurodevelopmental disorders. Copy number alterations (Sebat et al., 2007; Szatmari et al., 2007) and deleterious (Yan et al., 2008; Zahir et al., 2008) mutations in *αNrxn1* are the most commonly reported Nrxn isoform-specific modifications predisposing people to epilepsy, autism spectrum disorders (ASDs), attention deficit hyperactivity disorder, intellectual disability (ID), schizophrenia (SCZ) and Tourette syndrome (Ching et al., 2010; Clarke, Lee, & Eapen, 2012; Kim et al., 2008; Moller et al., 2013). Rare mutations in Nlgn3 have been reported in patients with ID, SCZ and ASDs (Jamain et al., 2003; Parente et al., 2017; Yan et al., 2005). Interestingly, ASD patients show abnormalities in memory discrimination (Beversdorf et al., 2000), which is partly mediated by the activity of hippocampal Cck+ interneurons (Sun et al., 2020; Whissell et al., 2019). This autistic phenotype may be caused by abnormal trans-synaptic interactions of αNrxn1 and Nlgn3 at VGT3+ synapses. In addition to our findings here, both Nrxn1 and Nlgn3 are expressed at other synapses, including excitatory synapses. (Baudouin et al., 2012; Budreck & Scheiffele, 2007; Uchigashima et al., 2019; Uchigashima et al., 2020) It will be interesting to identify Nrxn1-Nlgn3 codes at different synapses to better understand the function of these genes on cognitive behavior.

## Materials and Methods

### Animal and organotypic slice culture preparation

All animal protocols were approved by the Institutional Animal Care and Use Committee (IACUC) at the University of Massachusetts Medical School. Organotypic hippocampal slice cultures were prepared from postnatal 6- to 7-day-old mice of either sex, as described previously (Stoppini, Buchs, & Muller, 1991). Mice were wild-type (C57BL/6J, Jax #000664) expressing interneuron-specific TdTomato (Sst/RFP, Pv/RFP and VGT3/RFP), generated by crossing a TdTomato reporter line (Jax #007905) with Sst-Cre (Jax #013044), Pv-Cre (Jax #017320 or #008069) or VGT3-Cre (Jax #018147) lines. The Nrxn1/2/3^f/f^ mouse line was generated recently (Uemura et al., 2020).

### DNA and shRNA constructs

Enhanced green fluorescent protein (EGFP, Clontech), Tag-blue fluorescent protein (Tag-BFP, Evrogen), *Nlgn3Δ, αNrxn1±AS4, αNrxn3±AS4* and *βNrxn3±AS4* genes were subcloned into a pCAG vector. The full AS configuration of the *+Nrxn1* and *αNrxn3* clones were *αNrxn1(+AS1, -AS2, +AS3, ±AS4, +AS5* and *-AS6)* and *αNrxn3(-AS1, -AS2, +AS3, ±AS4, -AS5* and *-AS6)*. Two shRNA clones against Nlgn3, shNlgn3#1 (TRCN0000031940) and shNlgn3#2 (TRCN0000031939), were obtained from the RNAi Consortium (http://www.broad.mit.edu/genome_bio/trc/). The mouse *αNrxn3* clone was a gift from Dr. Ann Marie Craig.

### Antibodies

Primary antibodies raised against the following molecules were used: goat VGT3 (AB_2571854) (J. Somogyi et al., 2004), rabbit and goat VIAAT (RRID : AB_2571623 and AB_2571622) (Fukudome et al., 2004), rabbit CB1 (RRID : AB_2571591) (Fukudome et al., 2004), guinea pig Nlgn3 (RRID : AB_2571814) (Uchigashima et al., 2019; Uchigashima et al., 2020) rabbit PV (RRID : AB_2571613, Sigma: P3088) (Nakamura et al., 2004), rabbit Sst (Penisula lab.: T-4103.0050), and goat/rabbit RFP (Rockland: 200-101-379 and 600-401-379, respectively).

### Single- and double-labeled fluorescent *in situ* hybridization

Single/double FISH was performed using our recently established protocol (Uchigashima et al., 2019). All procedures were performed at room temperature unless otherwise noted. Briefly, fresh frozen sections were fixed with 4% paraformaldehyde, 0.1M PB for 30 min, acetylated with 0.25% acetic anhydride in 0.1M triethanolamine-HCl (pH 8.0) for 10 min, and prehybridized with hybridization buffer for 30 min. Hybridization was performed with a mixture of fluorescein- (1:1000) or DIG- (1:10,000) labeled cRNA probes in hybridization buffer overnight followed by post-hybridization washing using saline-sodium citrate buffers at 75°C. Signals were visualized using a two-step detection method. Sections were pre-treated with DIG blocking solution for 30 min and 0.5% tryamide signal amplification (TSA) blocking reagent in Tris-NaCl-Tween 20 (TNT) buffer for 30 min before the first and second steps. During the first step, sections were incubated with peroxidase-conjugated anti-fluorescein antibody (1:500, Roche Diagnostics) and TSA Plus Fluorescein amplification kit (PerkinElmer) for 1 hour and 10 min, respectively. In the second step, sections were incubated with peroxidase-conjugated anti-DIG antibody (1:500, Roche Diagnostics) and TSA Plus Cy3 amplification kit (second step) with the same incubation times. Between the first and second steps, residual peroxidase activity was inactivated with 3% H_2_O_2_ in TNT buffer for 30 min. Sections were incubated with DAPI for 10 min for nuclear counterstaining (1:5000, Sigma-Aldrich).

### Immunohistochemistry

#### Validation of VGT3/RFP, Sst/RFP, PV/RFP mouse lines

PFA-fixed brains (two brains for each line) were sliced (40 μm) and subjected to double staining with RFP (Rockland 200-101-379 or 600-401-379: 1/2000) and VGT3 (1 μg/ml) (J. Somogyi et al., 2004), Sst (Penisula lab., T-4103.0050: 1/2000) or PV (Sigma: P3088, 1/2000).

#### Triple staining of Nlgn3 and synaptic markers

Mice were fixed by transcardial perfusion with 3% glyoxal fixative (Richter et al., 2018). Brains were cryoprotected with 30% sucrose in 0.1 M PB to prepare 50 μm-thick cryosections on a cryostat (CM1900; Leica Microsystems). All immunohistochemical incubations were performed at room temperature. Sections were incubated with 10% normal donkey serum for 20 min, a mixture of primary antibodies overnight (1 μg/ml), and a mixture of Alexa 488-, Cy3-, or Alexa 647-labeled species-specific secondary antibodies for 2 hr at a dilution of 1:200 (Invitrogen; Jackson ImmunoResearch, West Grove, PA). Images were taken with a confocal laser scanning microscope equipped with 473, 559 and 647 nm diode laser lines, and UPlanSApo (10×/0.40), UPlanSApo (20×/0.75) and PlanApoN (60×/1.4, oil immersion) objective lenses (FV1200; Olympus, Tokyo, Japan). To avoid crosstalk between multiple fluorophores, Alexa488, Cy3, and Alexa647 fluorescent signals were acquired sequentially using the 488 nm, 543 nm and 633 nm excitation laser lines. All images show single optical sections (800 × 800 pixels). The signal intensity of Nlgn3 labeling in the hippocampus was measured using the Integrated Morphometry Analysis (IMA) function in MetaMorph software (Molecular Devices). Briefly, images were separated into individual channels, converted into grayscale, and thresholded based on CB1, VGT3, PV or Sst signals. The region of interest (ROI) for CB1, VGT3, PV or Sst punctum was created using Create Regions Around Objects function. Nlgn3 signal intensity was measured in CB1-, VGT3-, PV- or Sst-defined ROI.

### Single-cell sequencing and analysis

#### Single-Cell RNA Extraction

The cytosol of four VGT3-positive neurons in CA1 origins was harvested using the whole-cell patch-clamp technique described previously (Futai et al., 2013; Uchigashima et al., 2020). The cDNA libraries were prepared using a SMART-Seq^®^ HT Kit (TAKARA Bio) and a Nextera XT DNA Library Prep Kit (Illumina) as per the manufacturers’ instructions. The final product was assessed for its size distribution and concentration using a BioAnalyzer High Sensitivity DNA Kit (Agilent Technologies) and loaded onto an S1 flow cell on an Illumina NovaSeq 6000 (Illumina) and run for 2 x 50 cycles according to the manufacturer’s instructions. De-multiplexed and filtered reads were aligned to the mouse reference genome (GRCm38) using HISAT2 (version 2.1.0) (https://genomebiology.biomedcentral.com/articles/10.1186/gb-2013-14-4-r36) applying --no-mixed and --no-discordant options. Read counts were calculated using HTSeq (http://bioinformatics.oxfordjournals.org/content/31/2/166) by supplementing Ensembl gene annotation (GRCm38.78). Gene expression values were calculated as transcripts per million (TPM) using custom R scripts. Genes with no detected TPM in all samples were filtered out. The log2+1 transformed TPM values were combined with “Cell Diversity in the Mouse Cortex and Hippocampus RNA-Seq Data” from the Allen Institute for Brain Science (https://portal.brain-map.org/atlases-and-data/rnaseq#Mouse_Cortex_and_Hip) where large-scale single-cell RNA-seq data were collected from adult mouse brain. RNA sequencing data were generated from single cells isolated from >20 areas of mouse cortex and hippocampus, including ACA, AI, AUD, CA, CLA, CLA;EPd, ENTl, ENTm, GU;VISC;AIp, HIP, MOp, MOs, ORB, PAR;POST;PRE, PL;ILA, PTLp, RSP, RSPv, SSp, SSs, SSs;GU, SSs;GU;VISC, SUB;ProS, TEa;PERI;ECT, VISal;VISl;VISli, VISam;VISpm, VISp and VISpl;VISpor (Abbreviations for each cell type match those found in the Allen Mouse Brain Atlas (https://portal.brain-map.org/explore/classes/nomenclature)). The provided table of median expression values for each gene in each cell type cluster was merged with our data, and a tSNE plot was generated using Rtsne R package (van der Maaten and Hinton, 2008). For splice isoform quantification, kallisto (Bray et al., 2016) was used by supplementing the transcript fasta file (Mus_musculus.GRCm38.cdna.all.fa). Each isoform was summarized manually to account for inclusion or exclusion of the AS4 exon in the α or β isoforms. The manually curated transcript IDs are provided in **Table 1**.

### Single-cell RT-qPCR

Isolation of single-cell cytosol and preparation of single-cell cDNA libraries were performed by the same method described in **Single-cell sequencing and analysis**. For validation of the Nrxn KO mouse line, the following TaqMan gene expression assays (Applied Biosystems) were used: *Nrxn1* (Mm03808857_m1), *Nrxn2* (Mm01236856_m1), *Nrxn3* (Mm00553213_m1), *Slc17a8* (VGT3: Mm00805413_m1) and *Gapdh* (Mm99999915_g1). The relative expression of *Nrxns* or *Slc17as* were calculated as follows: Relative expression = 2^Ct,Gapdh^/2 ^Ct,Nrxns or Slc17as^; Ct, threshold cycle for target gene amplification.

### Single-cell electroporation

A detailed protocol is described in our recent publication (Keener et al., 2020). Briefly, the slice cultures were perfused with filter-sterilized aCSF consisting of (in mM): 119 NaCl, 2.5 KCl, 0.5 CaCl_2_, 5 MgCl_2_, 26 NaHCO_3_, 1 NaH_2_PO_4_, 11 glucose and 0.001 mM tetrodotoxin (TTX, Hello Bio Inc), gassed with 5% CO_2_/95% O_2_, pH 7.4. Patch pipettes (4.5 – 8.0 MΩ) were each filled with plasmids containing either tag-BFP and *Nrxn* or EGFP and *Nlgn3Δ* (0.05 μg/μl for each plasmid) and respectively electroporated in TdTomato-positive VGT3+ interneurons and CA1 pyramidal neurons. The same internal solution for single-cell sequencing was used. A single electrical pulse train (amplitude: −5 V, square pulse, train: 500 ms, frequency: 50 Hz, pulse width: 500 μs) was applied to the target neurons. After electroporation, the electrode was gently retracted from the cell and the slices were transferred to a culture insert (Millipore) with slice culture medium in a petri dish and incubated in a 5% CO_2_ incubator at 35°C for 3 days.

### Electrophysiology

Whole-cell voltage- and current-clamp recordings were performed on postsynaptic and presynaptic neurons, respectively. *Nlgn3Δ* construct or shRNAs were transfected at DIV6-9 or DIV2 and subjected to recordings at two to three or five to twelve days after transfection, respectively. The extracellular solution for recording consisted of (in mM): 119 NaCl, 2.5 KCl, 4 CaCl_2_, 4 MgCl_2_, 26 NaHCO_3_, 1 NaH_2_PO_4_, 11 glucose and 10 kynurenic acid (Sigma), gassed with 5% CO_2_ and 95% O_2_, pH 7.4. Thick-walled borosilicate glass pipettes were pulled to a resistance of 2.5 – 4.5 MΩ. Whole-cell voltage clamp recordings were performed with internal solution containing (in mM): 115 cesium methanesulfonate, 20 CsCl, 10 HEPES, 2.5 MgCl_2_, 4 ATP disodium salt, 0.4 guanosine triphosphate trisodium salt, 10 sodium phosphocreatine, and 0.6 EGTA, adjusted to pH 7.25 with CsOH. For current-clamp recordings, cesium in the internal solution was substituted with potassium and the pH was adjusted with KOH. GABAA receptor-mediated inhibitory postsynaptic currents (IPSCs) were measured at *V*hold ± 0 or −70 mV. Thirty to forty consecutive stable postsynaptic currents were evoked at 0.1 Hz by injecting current (1 nA) in presynaptic interneurons. Recordings were performed using a MultiClamp 700B amplifier and Digidata 1440, digitized at 10 kHz and filtered at 4 kHz with a low-pass filter. Data were acquired and analyzed using pClamp (Molecular Devices).

### Statistical analyses

Results are reported as mean ± SEM. Statistical significance, set at p < 0.05, was evaluated by two-way ANOVA with *post hoc* Tukey for multiple comparison, Mann-Whitney U-test and Student’s t-test for two-group comparison.

### DATA availability

The accession number for the RNA-seq and processed data reported in this paper is GEO: GSE150989.

## Acknowledgments

This work was supported by grants from the National Institutes of Health Grants (R01NS085215 to K.F., T32 GM107000 and F30MH122146 to A.C.), the Global Collaborative Research Project of Brain Research Institute, Niigata University (G2905 to K.F.), Grants-in-Aid for Scientific Research (19100005 to M.W.; 15K06732 and 20H03349 to M.U.) and The Naito Foundation (to M.U.). The authors thank Ms. Naoe Watanabe and Ms. Rie Natsume for skillful technical assistance. We thank Dr. Paul Gardner for comments on an earlier draft of the manuscript.

## Author contributions

M.U. and K.F. designed this research. M.U., K.K., E.D., A.C., T.W., D.K., M.A., T.L., T.U., Y.I.K., and K.F. performed the experiments and analyzed the data. M.U., K.K., A.C., T.W., K.S., T.S., T.U., Y.I.K., M.W, and K.F. wrote the paper.

---

**Figure S1.**
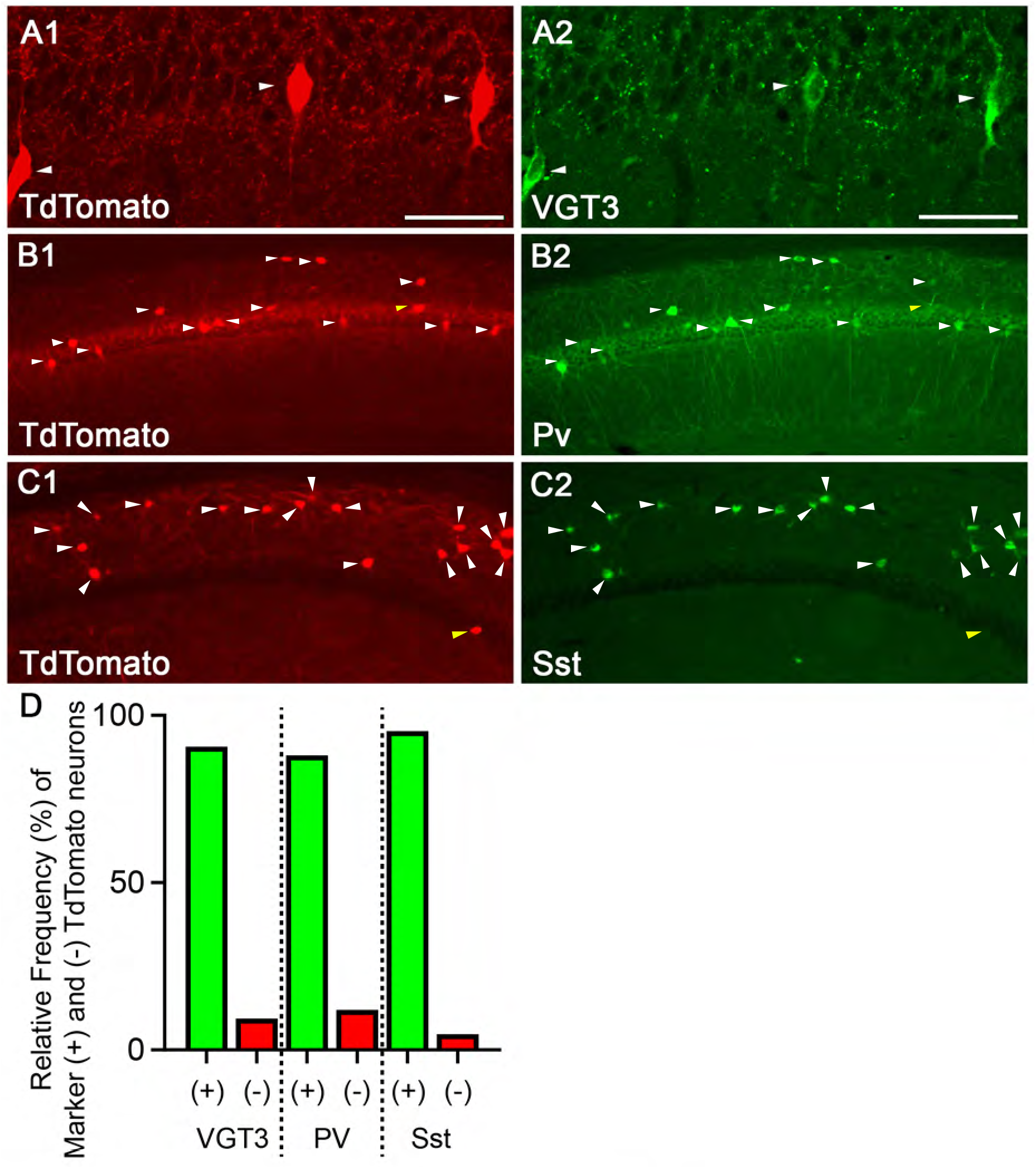
Validation of TdTomato expression in three different cell type-specific fluorescent lines. **(A-C)** Images of double-labeled immunofluorescence staining against TdTomato (red, **A1, B1, C1**) and cell type-specific markers (green, VGT3: **A2**, Pv: **B2**, Sst: **C2**) in the hippocampal CA1 region of VGT3/RFP **(A)**, Pv/RFP **(B)** and Sst/RFP **(C)** mice. **(D)** Relative proportion of hippocampal CA1 inhibitory interneurons as a fraction of neuronal populations positive for cell type-specific markers and TdTomato (+) and TdTomato-positive but marker negative (-) neurons. The TdTomato-positive neurons included for quantification were localized in the specific CA1 subregions where we performed electrophysiological recordings; VGT3/RFP mice: CA1 pyramidale, radiatum and oriens; Pv/RFP: radiatum; Sst/RFP: oriens. Number of TdTomato-positive neurons: VGT3/RFP: 404 neurons/2 mice; Pv/RFP: 1601/2; Sst/RFP: 1789/2. Scale bars, 50 μm.

**Figure S2.**
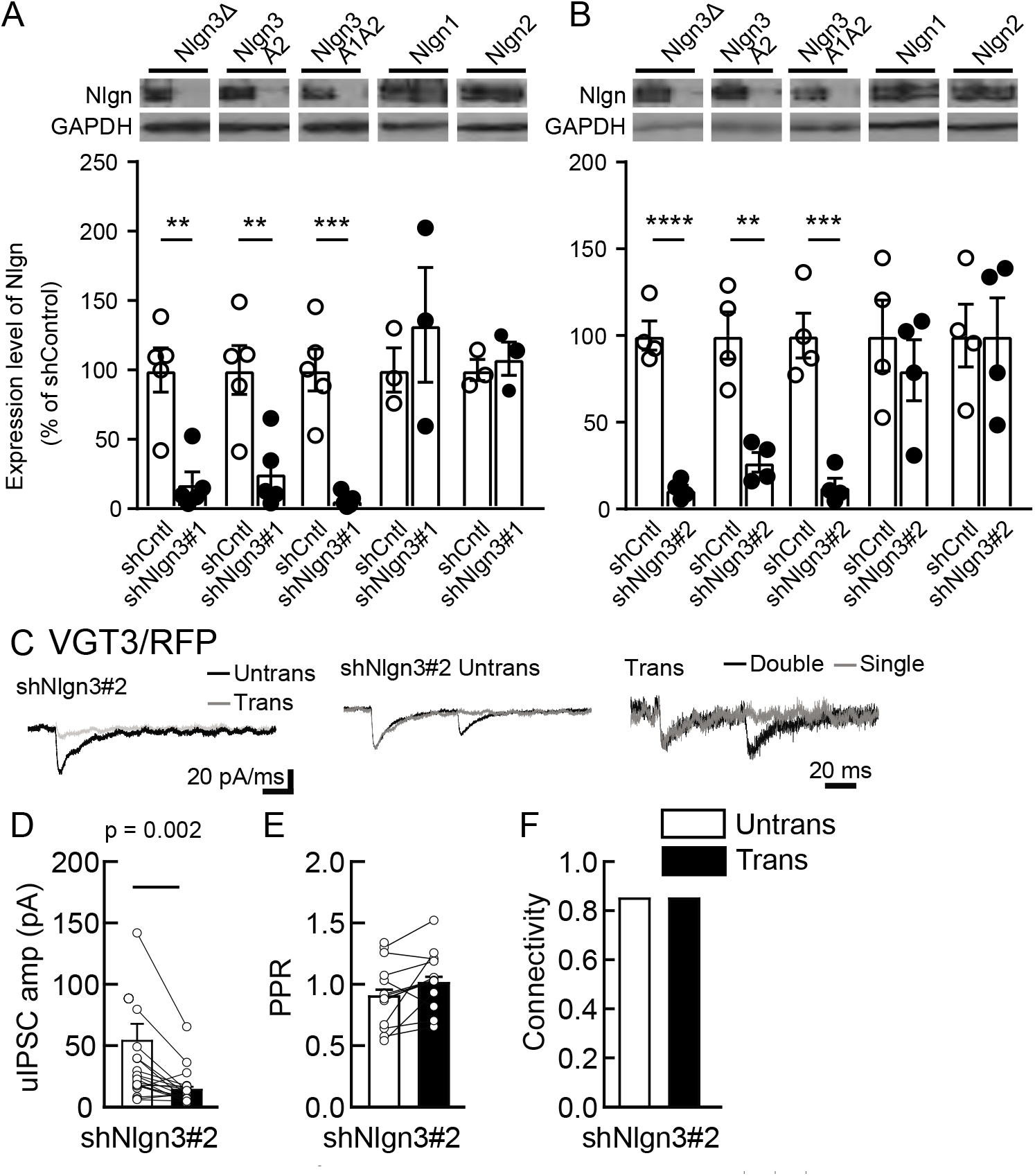
Nlgn3 shRNAs specifically knockdown Nlgn3 protein and regulate VGT3+ inhibitory synaptic transmission. **(A, B)** Specificity of Nlgn3 shRNA constructs, shNlgn3#1 **(A)** and shNlgn3#2 **(B)**, confirmed in HEK293T cell lysates overexpressing HA-tagged Nlgn3Δ, Nlgn3A2, Nlgn3A1A2, Nlgn1 and Nlgn2. (Top) Immunoblot images with HA and GAPDH. Membranes were first probed with rabbit HA antibody, stripped, then re-probed with mouse GAPDH antibody. (Bottom) Summary bar graphs with individual scattered plots of shRNA effects. **(C-F)** Effects of shNlgn3#2 expression in hippocampal CA1 pyramidal neurons were compared between three different inhibitory inputs mediated by VGT3+ inhibitory interneurons. **(C)** Sample traces of uIPSCs measured at −70 mV. Left, superimposed averaged sample traces of uIPSC (Untrans: black, trans: dark gray) induced by an AP. Middle and right, superimposed averaged sample traces of uIPSCs evoked by single (dark gray) and double (black) APs in VGT3+ interneurons. uIPSCs are normalized to the first amplitude. The amplitude **(D)** and PPR **(E)** of uIPSCs were plotted for each pair of transfected (Trans) and neighboring untransfected (Untrans) cells (open symbols). Bar graphs indicate the mean ± SEM. **(F)** Synaptic connectivity between presynaptic inhibitory interneuron and postsynaptic untransfected (open bars) or transfected (black) pyramidal neurons. Numbers of cell pairs: shNlgn3 at VGT3+ synapses (18 cell pairs/10 mice). The number of tested slice cultures is the same as that of cell pairs. n.s., not significant. Mann–Whitney U test.

**Figure S3.**
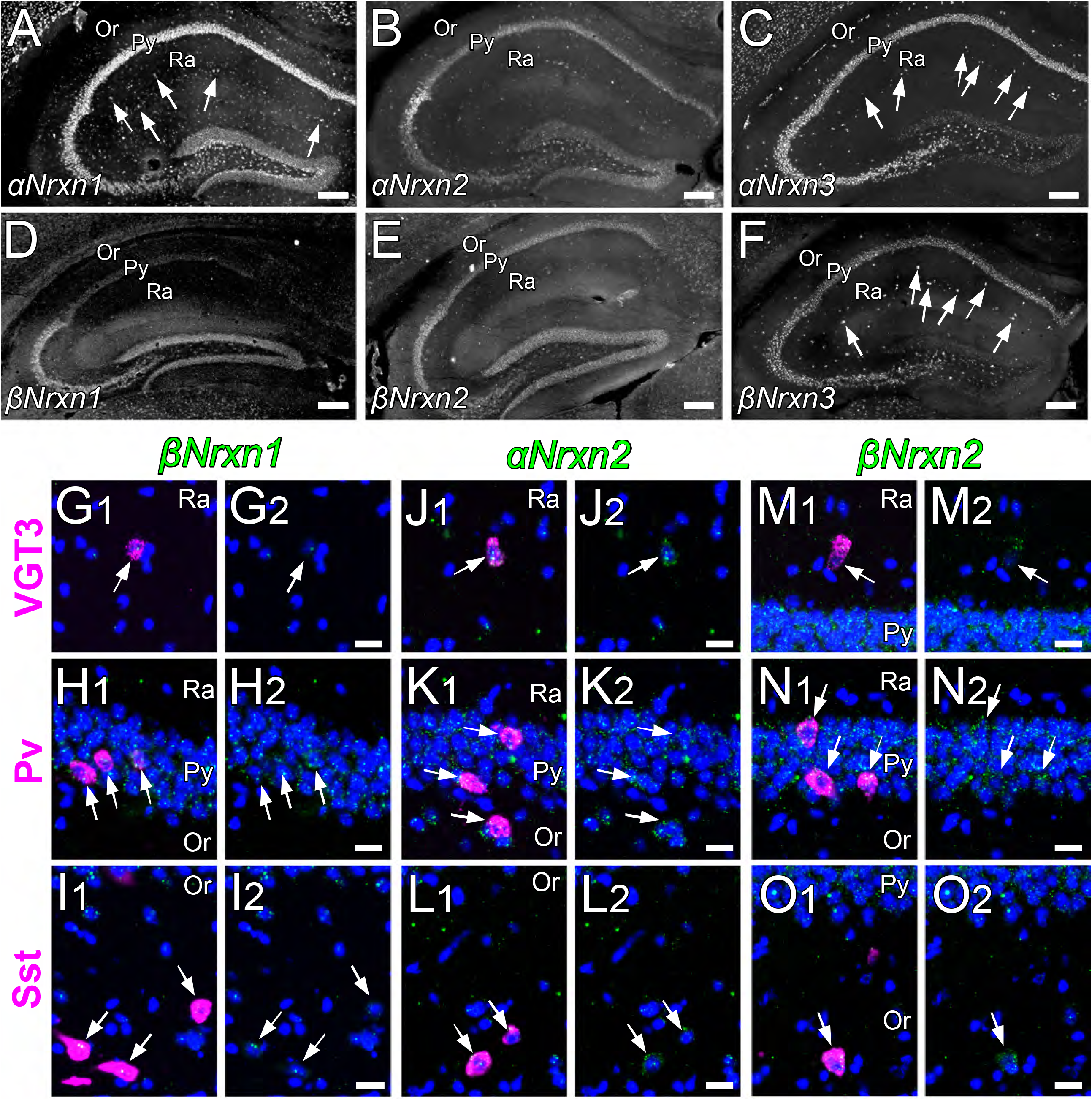
Expression of *Nrxn* isoforms in different types of inhibitory interneurons in the hippocampal CA1 region. **(A-F)** Single FISH for *αNrxn1* **(A)**, *αNrxn2* **(B)**, *αNrxn3* **(C)** and *βNrxn1* **(D)**, *βNrxn2* **(E)** and *βNrxn3* **(F)** in the hippocampus. Striking signals for *αNrxn1* **(A)**, *αNrxn3* (**C**) and *βNrxn3* **(F)** are noted in sparse cells in the st. radiatum (arrows). **(G-O)** Double FISH for *βNrxn1* **(G, H, I)**, *αNrxn2* **(J, K, L)** and *βNrxn2* **(M, N, O)** in VGT3+ **(G, J, M)**, Pv+ **(H, K, N)** and Sst+ **(I, L, O)** in the hippocampus showing different levels of *Nrxn* mRNA (green) expression in different inhibitory interneurons (magenta, arrows). Note that the signal intensity in individual GABAergic neurons is variable, compared with that in glutamatergic neurons. Nuclei were stained with DAPI (blue). Or, st. oriens; Py, pyramidal cell layer; Ra, st. radiatum. Note that none of the Nrxn isoforms displayed unique expression specific to VGT3+ inhibitory interneurons (See **Figure 6J**). Scale bars, 200 μm **(A-F)**, 20 μm **(G-O)**.

**Figure S4.**
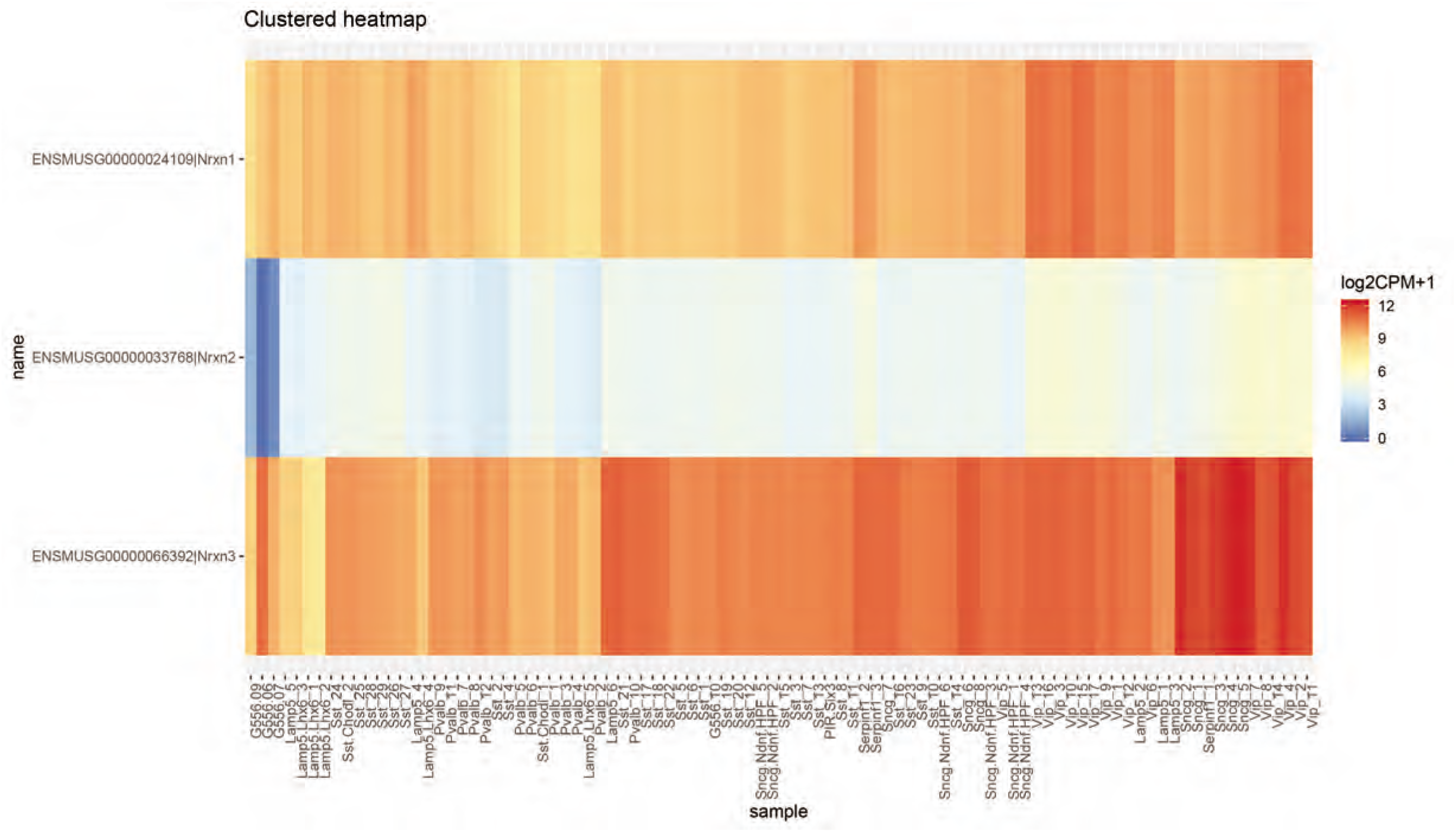
Heat map of *Nrxn* gene expression in VGT3+ inhibitory interneurons. A hierarchical clustering of Nrxn expression levels demonstrated similarities between the G556 cells and adult GABAergic neurons, which clustered aside from the GABAergic neurons potentially due to relatively lower expression of Nrxn2 in the G556 cells. Scale bar shows log2+1 transformed CPM values.

